# Large Scale Discovery of Microbial Fibrillar Adhesins and Identification of Novel Members of Adhesive Domain Families

**DOI:** 10.1101/2021.12.07.471604

**Authors:** Vivian Monzon, Alex Bateman

**Affiliations:** European Molecular Biology Laboratory, European Bioinformatics Institute (EMBL-EBI), Wellcome Genome Campus, Hinxton. CB10 1SD

**Keywords:** Fibrillar adhesins, Host-pathogen interaction, RandomForest classification, Protein domain families, Adhesive domains, Structure prediction methods, AlphaFold2

## Abstract

Fibrillar adhesins are bacterial cell surface proteins that mediate interactions with the environment including host cells during colonisation or other bacteria during biofilm formation. These proteins are characterised by a stalk that projects the adhesive domain closer to the binding target. Fibrillar adhesins evolve quickly and thus can be difficult to computationally identify, yet they represent an important component for understanding bacterial host interactions.

To detect novel fibrillar adhesins we developed a random forest prediction approach based on common characteristics we identified for this protein class. We applied this approach to Firmicute and Actinobacterial proteomes, yielding over 6,500 confidently predicted fibrillar adhesins. To verify the approach we investigated predicted fibrillar adhesins that lacked a known adhesive domain. Based on these proteins, we identified 24 sequence clusters representing potential novel members of adhesive domain families. We used AlphaFold to verify that 15 clusters showed structural similarity to known adhesive domains such as the TED domain.

Overall our study has made a significant contribution to the number of known fibrillar adhesins and has enabled us to identify novel members of adhesive domain families involved in the bacterial pathogenesis.

**Importance:** Fibrillar adhesins are a class of bacterial cell surface proteins that enable bacteria to interact with their environment. We developed a Machine Learning approach to identify fibrillar adhesins and applied this classification approach on the Firmicutes and Actinobacteria Reference Proteomes. This method allowed us to detect a high number of novel fibrillar adhesins, and also novel members of adhesive domain families. To confirm our predictions of these potential adhesin protein domains, we predicted their structure using the AlphaFold tool.

## Introduction

Fibrillar adhesins are an important class of bacterial surface proteins, which are expressed by a wide range of bacterial species to mediate binding interactions. Essential binding targets include different host surface structures, such as extracellular matrix proteins. A pathogenic colonization and infection can occur as a consequence of the binding interactions to host cells [1]. Single fibrillar adhesins have therefore been studied in depth and have been the focus point for anti-adhesion therapies [2, 3]. Fibrillar adhesins have also been described to mediate biofilm formation [4, 5].

Fibrillar adhesins are a recently defined class of proteins that led to a domain-based characterization and identification of these proteins across a wide range of bacterial species [6, 7]. Key characteristics of fibrillar adhesins are their large length with an adhesion region, a rod-like repetitive region and a cell surface anchor. The repetitive region contains repeating protein domains, also called stalk domains, which fold into a filamentous stalk. It has been suggested that the stalk projects the binding region closer to the binding target [8] and enables the adhesive region to be presented outside the cell by reaching beyond the surface layer. The repeats can vary in number leading to fibrillar adhesins with stalks of different lengths. Proteins with varying repeat number of stalk domains between related bacterial strains have recently been termed ‘Periscope proteins’ [9]. Whelan *et al.* propose that the varying length is used as a regulation mechanism to facilitate the binding to targets at different distances [9]. The varying length may also lead to the adhesive region being differentially displayed beyond the surface layer based on the number of repeats. Several adhesive domains have been identified when studying the binding regions of bacterial adhesins, of which most bind to protein ligands, some bind to carbohydrates and one adhesive domain is known to bind to ice crystals [10]. Nevertheless, undoubtedly a large number of adhesive domains remain undiscovered.

In our previous work, we have used the presence of known adhesive and stalk domains identified in fibrillar adhesins to detect more than 3,000 fibrillar adhesins-like (FA-like) proteins across all bacterial species based on profile hidden Markov models (HMM)-searches [7]. The limitation of this domain-based discovery approach is that only FA-like proteins with known adhesive and stalk domains are found. Not all adhesive and stalk domains are identified yet and finding novel binding proteins or domains is important for the understanding of emerging bacterial interactions. To overcome this limitation we studied the properties of FA-like proteins and developed a random-forest based discovery approach. The aim of this study is to enable the identification of FA-like proteins on a large-scale, including those lacking a known adhesive or stalk domain. We apply our newly developed machine learning approach on the Firmicute and Actinobacteria UniProt Reference Proteomes and verify the approach by predicting the structure of Firmicute FA-like proteins lacking a known adhesive domain with AlphaFold [11]. Our approach facilitates the identification of relevant proteins during bacterial infection processes, enabling the investigation of novel members of adhesive domain families leading to a better understanding of microbial interaction mechanisms.

## Results

### Random Forest classification

The aim of this study is the extension of the identification of fibrillar adhesins compared to the domain-based approach described in our earlier study [7]. To achieve this goal, additional identification features were combined with the presence of adhesive and stalk domains, although the adhesive and stalk domains clearly remain the most important features in this study. To undertake the identification approach based on the selected features a Random Forest approach was selected. This is composed of individual decision trees classifying the proteins by sets of maximum three features of all identification features provided as input to the algorithm and returning the strongest class as prediction.

We decided to concentrate on the Firmicute and Actinobacteria phyla in this study. Fibrillar adhesins are best studied in Firmicutes and the cell surface composition and FA-like protein architecture of the Actinobacteria resembles those of the Firmicutes [3, 7]. We created a positive training set based on the FA-like proteins identified in our previous study [7]. For the negative training set we randomly selected non-FA-like proteins from eight Firmicute and Actinobacteria Reference Proteomes, in which FA-like proteins were detected in our previous study [7]. Although an individual proteome may contain several FA-like proteins, they are present at relatively low numbers per proteome (0% to 1.47%), thus randomly selecting the negative examples is unlikely to select many, if any, true FA-like proteins [7]. In total, the training set consists of 3,332 proteins, equally balanced between the positive and negative training sets. Using this training set, common properties for this protein class were determined.

Nearly all proteins of the positive training set (98%) have at least one adhesive and one stalk domain, whereas less than 0.1% of the proteins in the negative training set possess a known adhesive or stalk domain. Hence, the adhesive and stalk domains are the strongest identification features for this protein class (Figure 1). To increase the chance to detect proteins with unidentified stalk domains, we selected the presence of tandem sequence repeats with a minimum of 70% sequence identity allowed by the T-REKS tool [12]. In periscope proteins the stalk domains are described to be found in highly identical tandem repeats [9]. But compared to periscope proteins, not all stalk domains in FA-like proteins are found in highly identical tandem repeats and are therefore missed by T-REKS. We observed that the stalk domain region tends to have a biased sequence composition and tends to be predicted as disordered despite known structures being found in some of these regions. We used the fraction of predicted disordered residues by IUPRED2, as an additional feature [13].

**Figure 1:**
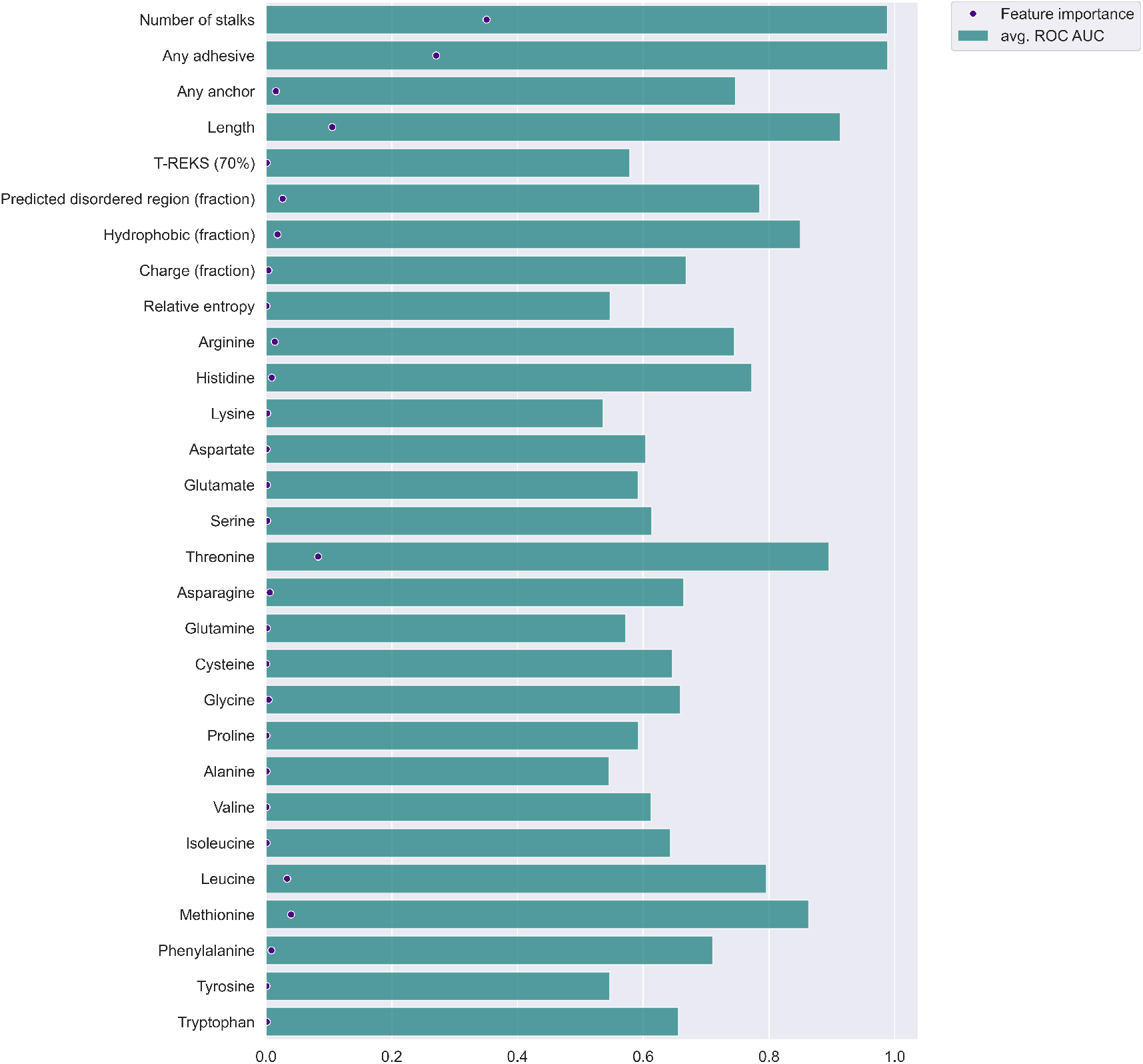
A bar plot showing the relative importance of prediction features: This plot visualizes the importance of each feature for the random forest classification. The bars show the calculated ROC AUC per feature when using it alone for classification. The dot represents the feature importance as calculated by the feature importance attribute implemented in the sklearn ensemble random forest classifier.

Fibrillar adhesins are attached to the bacterial cell surface. Known anchor domains or sortase motifs were found in 846 out of 1666 proteins of the positive training set and were used as another identification feature.

FA-like proteins are among the longest proteins in the protome, thus the protein length turned out to be one of the strongest prediction features (Figure 1). Their high length facilitates FA-like proteins to cross the peptidoglycan layer, which can be around 20-50 nm wide depending on the bacterial species [14]. The average protein length of the positive training set is 1,196 residues compared to 300 residues for the negative training set. Hence, with a higher sequence length of a protein, the probability increases that the protein functions as a FA-like protein.

To characterize the protein sequence of FA-like proteins, the amount of charged as well as hydrophobic amino acids per protein were selected as additional features. The protein sequences of the positive training data tend to have a slightly lower fraction of charged amino acids and tend to have a lower fraction of hydrophobic amino acids compared to the negative training data set.

Finally, we selected features related to the sequence composition. We calculated the fraction of each amino acid per sequence as well as the relative entropy describing the sequence composition bias. We observed that the relative entropy tends to be slightly higher in the positive training set and that threonine is 1.8-fold increased and leucine is 1.5-fold decreased in the positive training set compared to the negative training set (Figure S1).

We implemented the selected features in a random forest classification approach and analysed the feature importance in the classification prediction based on the training data (Figure 1). The adhesive and stalk domain features can yield a Receiver Operating Characteristics (ROC) Area Under the Curve (AUC) of 0.99 due to the fact that the training set is built upon FA-like proteins identified in our previous work using known adhesive and stalk domains (Figure 1). The low diversity of the training data, considering that the positive training data nearly solely consists of proteins with at least one adhesive and one stalk domain, is reflected in the reliability and precision-recall curve (Figure 2a,b). The reliability curve shows the high number of proteins of the negative training set predicted with a prediction score below 0.1 and the high number of proteins of the positive training set predicted with a score above 0.9. Even though the number of predicted proteins with a score between 0.2 and 0.8 are low, the ratio of false positives tends to increase in the prediction score ranges 0.5 to 0.7. To test if the training set leads to an overfitted model, we evaluated the random forest classification approach with an extreme case of FA-like proteins that we artificially present to have no adhesive or stalk domains (all other features of the proteins are retained), yielding a precision score of 0.8 and achieving a recall score of 0.67 due to missed true matches (Figure 2b).

**Figure 2:**
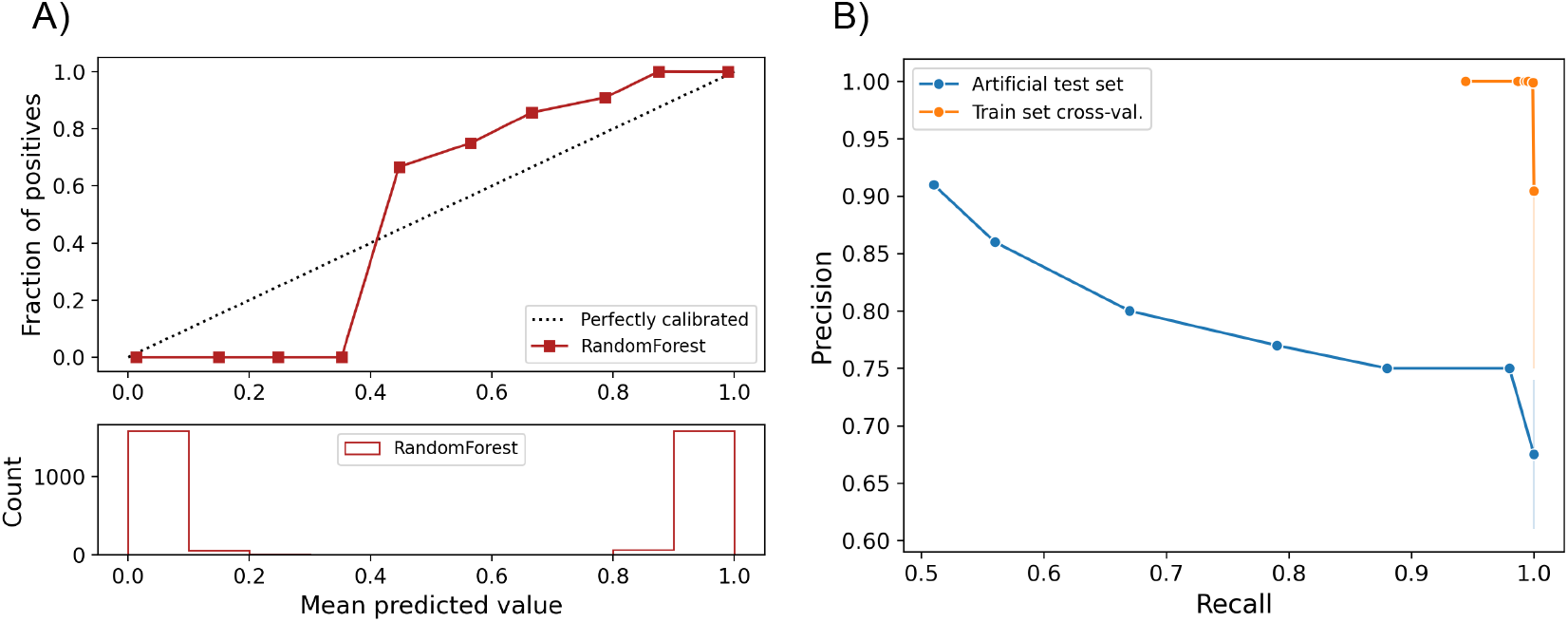
Validation of the trained random forest classifier: a) the subplot on the top represents the reliability curve showing the observed fraction of predicted proteins belonging to the positive training data set against the expected fraction of positives. The subplot below indicated the total number of proteins of the training set predicted per prediction score. b) Precision-Recall curves calculated with the training set using a cross-validation approach (orange) or using a test set with a positive set of FA-like proteins with adhesive and stalk domains artificially removed (blue).

An important challenge for this work is to determine the random forest score threshold that will reliably identify novel FA-like proteins that potentially lack known stalk or adhesive domains.

### Analysis of predicted FA-like proteins

When applying the classification approach on the Firmicute and Actinobacteria UniProt Reference Proteomes, 45,444 FA-like proteins with a prediction score above 0.5 were identified, 24,197 proteins in Firmicutes and 21,247 proteins in Actinobacteria (Table 1). These represent 0.49% and 0.32% of the total number of reference proteins respectively. The reference proteomes with the highest fraction of predicted FA-like proteins with a score of 0.7 or above are listed in supplementary table S1. Here we provide an analysis to help to determine a reasonable threshold to apply for downstream analysis and application of the classifier in general.

**Table 1:** Number of FA-like proteins discovered in Firmicutes and Actinobacteria per prediction score bin.

| Score | 0.5 – 0.6 | 0.6 – 0.7 | 0.7 – 0.8 | 0.8 – 0.9 | 0.9 – 1.0 | 1.0 |
| --- | --- | --- | --- | --- | --- | --- |
| Firmicutes | 6,325 | 7,110 | 5,751 | 2,694 | 1,130 | 1,187 |
| Actinobacteria | 6,715 | 8,718 | 4,121 | 924 | 303 | 466 |

To study the characteristics of the predicted FA-like proteins (at a threshold >0.5), we predicted their subcellular localizations using PSORTb (version 3) [15]. For nearly half of the predicted FA-like proteins in Firmicutes with a prediction score of 0.7 or below no localization was predicted, whereas the majority of FA-like proteins with a prediction score above 0.7 are predicted to be localised at the cell wall (Figure 3a). For most of the predicted FA-like proteins in Actinobacteria no localization was predicted by PSORTb. The highest protein number with a predicted localization are said to be extracellular, with around half of the FA-like proteins with a prediction score between 0.8 and 0.9 being predicted to be localised at the cell wall (Figure 3b). Consequently, the cell wall anchor characteristic of FA-like proteins is more strongly represented in the predicted proteins in Firmicutes compared to Actinobacteria.

**Figure 3:**
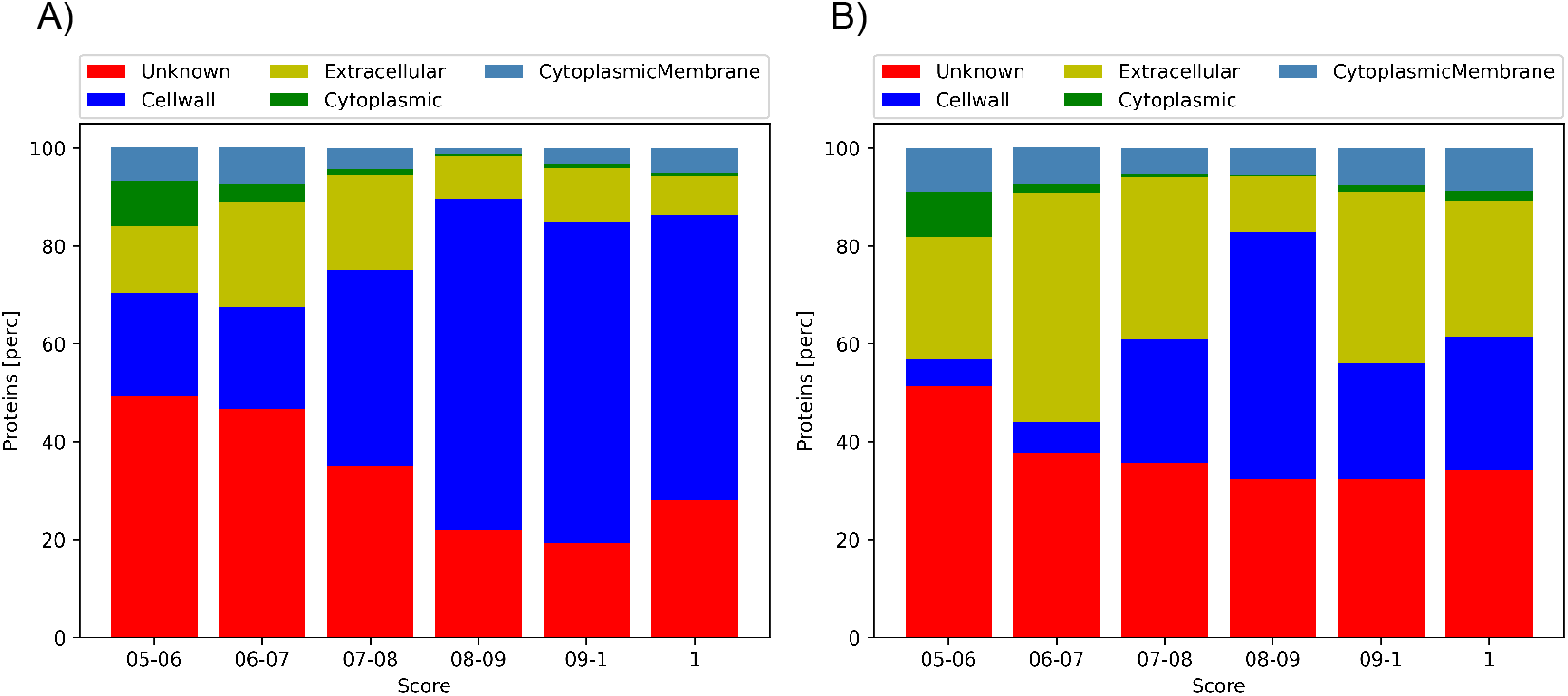
Predicted Subcellular Localization: By PSORTb predicted subcellular localizations for FA-like proteins predicted with a prediction score between 0.5 and 1.0 for a) Firmicutes and b) Actinobactria.

Even though most of the proteins (>75%) predicted with a score of 1.0 with known stalk domains have an adhesive domain, over 90% of the predicted FA-like proteins above a prediction score of 0.5 with known stalk domains have no known adhesive domain (Table 2a,b). Given that our training set is biased towards FA-like proteins with both stalk and adhesive domains, this result may suggest that there exist a large number of FA-like proteins with as yet undiscovered or unannotated adhesive domains.

**Table 2:**
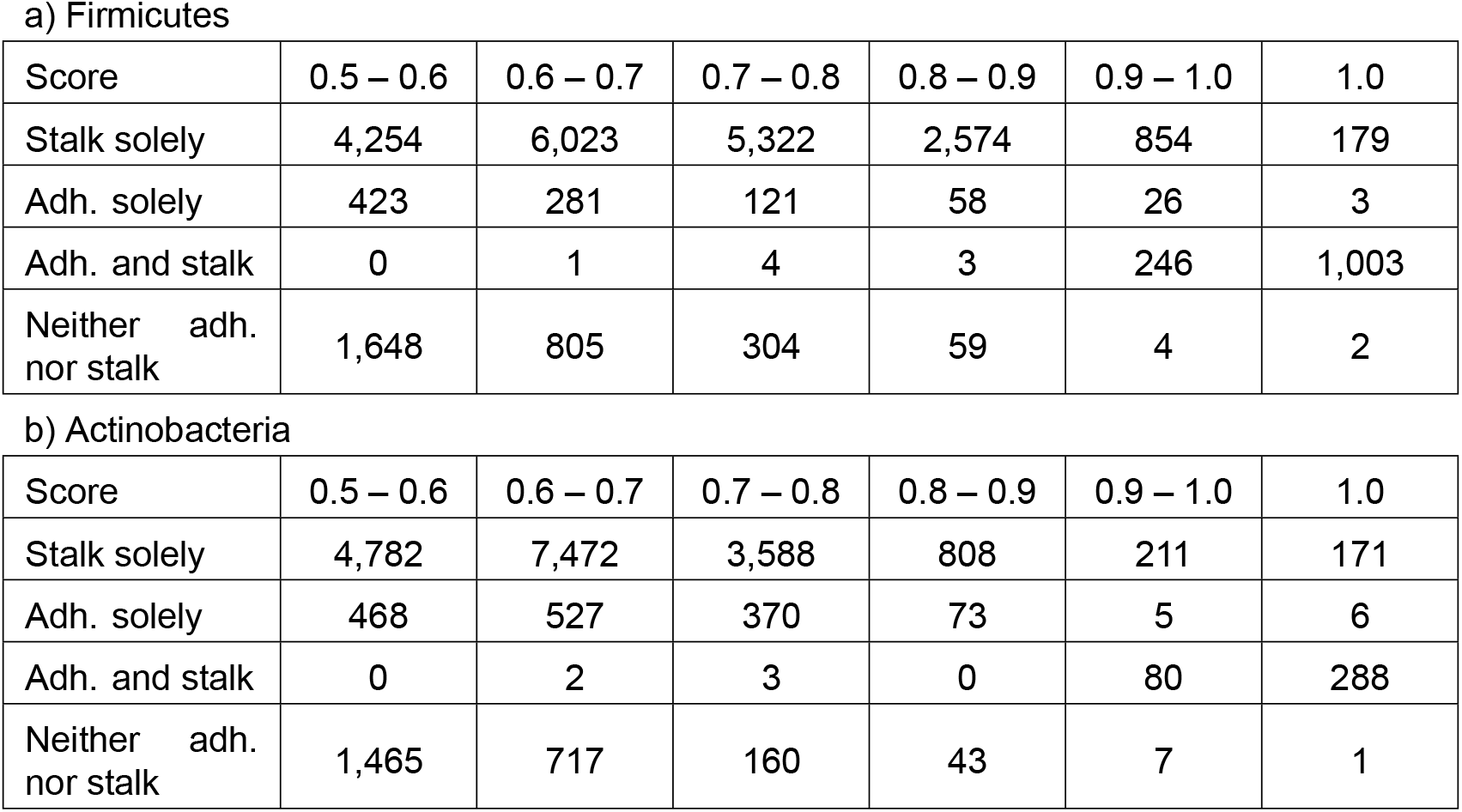
Overview of the presence and absence of known adhesive and stalk domains: The number of proteins per prediction score category is counted, differentiating between proteins with an adhesive and/or stalk domains or neither an adhesive nor a stalk domain. The results are listed for the predicted FA-like proteins in a) Firmicutes and b) Actinobacteria.

| Score | 0.5 – 0.6 | 0.6 – 0.7 | 0.7 – 0.8 | 0.8 – 0.9 | 0.9 – 1.0 | 1.0 |
| --- | --- | --- | --- | --- | --- | --- |
| Stalk solely | 4,254 | 6,023 | 5,322 | 2,574 | 854 | 179 |
| Adh. solely | 423 | 281 | 121 | 58 | 26 | 3 |
| Adh. and stalk | 0 | 1 | 4 | 3 | 246 | 1,003 |
| Neither adh.<br>nor stalk | 1,648 | 805 | 304 | 59 | 4 | 2 |

b) Actinobacteria
| Score | 0.5 – 0.6 | 0.6 – 0.7 | 0.7 – 0.8 | 0.8 – 0.9 | 0.9 – 1.0 | 1.0 |
| --- | --- | --- | --- | --- | --- | --- |
| Stalk solely | 4,782 | 7,472 | 3,588 | 808 | 211 | 171 |
| Adh. solely | 468 | 527 | 370 | 73 | 5 | 6 |
| Adh. and stalk | 0 | 2 | 3 | 0 | 80 | 288 |
| Neither adh.<br>nor stalk | 1,465 | 717 | 160 | 43 | 7 | 1 |

We investigated the predicted proteins missing an adhesive domain and found adhesive-domain-like sequences (Table 3). These were searched with the known adhesive domains using HMMER with a higher (less significant) E-value threshold of 1.0. Already over 60% of the proteins with a prediction score of 1.0 have a known adhesive domain (Table 3). Adding the number of proteins with distantly related adhesive domains leads to an increase up to 6.72% for a prediction score between 0.8 to 0.9 in Firmicutes (Table 3). These results suggest the presence of novel adhesive domain families distantly related to existing ones detected in the predicted FA-like proteins. The results also indicate the possible existence of potential novel adhesive domains, unrelated to known adhesive domains.

**Table 3:** Distantly related adhesive domains: Percentage of the total predicted FA-like proteins with only known adhesive domains and additional with the distantly related adhesive domains found by using a less significant Evalue.

| Score | 0.5 – 0.6 | 0.6 – 0.7 | 0.7 – 0.8 | 0.8 – 0.9 | 0.9 – 1.0 | 1.0 |
| --- | --- | --- | --- | --- | --- | --- |
| Firmicutes (only known adh.) | 6.69%<br>(423) | 3.97%<br>(282) | 2.17%<br>(125) | 2.26%<br>(61) | 24.07%<br>(272) | 84.75%<br>(1006) |
| Firmicutes (+ distantly related adh.) | 9.98%<br>(631) | 8.28%<br>(589) | 7.55%<br>(434) | 8.98%<br>(242) | 28.76%<br>(325) | 84.92%<br>(1008) |
| Actinobacteria (only known adh.) | 6.97%<br>(468) | 6.07%<br>(529) | 0.9%<br>(37) | 7.9%<br>(73) | 28.05%<br>(85) | 63.09%<br>(294) |
| Actinobacteria (+ distantly related adh.) | 7.55%<br>(507) | 7.66%<br>(668) | 3.45%<br>(142) | 10.71%<br>(99) | 28.38%<br>(86) | 63.09%<br>(294) |

Taking the observed results into account, we suggest a high confidence scoring threshold of 0.8, since the predicted FA-like proteins with a scoring threshold below 0.8 might include false positives. Nevertheless, a scoring threshold of 0.7 can be used to find an extended set of FA-like proteins as long as a careful verification of the predicted proteins is carried out.

### Verification of novel adhesive domains

To verify the random forest based discovery approach we further investigated the predicted FA-like proteins in the Firmicute Reference Proteomes. These include proteins without known adhesive or stalk domains. Here, we are interested in the proteins with known stalk domains that lack a known adhesive domain. We observed that many of them have a domain annotation gap at the N-terminus, distal to the cell surface anchor. Of these proteins we selected a subset of proteins with a minimum of 4 stalk domains. These were 1,546 proteins in Firmicutes with a prediction score above 0.5. Under the assumption that these annotation gaps might include an adhesive region of the proteins, we clustered the N-terminal sequences into homologous sequence clusters using blastp [16]. We selected the clusters with more than 5 sequences and with an average prediction score of 0.7 or above for further investigation, resulting in 24 clusters. To further investigate the clusters we chose one representative sequence per cluster (Table S2). We carried out two analyses (i) to search for overlapping known Pfam domains, using the highly sensitive iterative Jackhmmer search, and (ii) to predict the structure of the sequence using AlphaFold [17, 11]. Using Jackhmmer can be considered a more sensitive version of our previous analysis where we used a less significant HMMER threshold for known Pfam adhesive domains. Using Jackhmmer we are able to detect even more distant similarities to known domains. To find out more about the function, particularly for sequences without overlapping Pfam domains, we searched with the predicted AlphaFold structure models against the PDB database for similar structures [18] (Suppl. Table S3). We aligned the sequences of each cluster and built profile-HMMs specific to each cluster. To understand the relative abundance of each of our clusters we then searched for homologous sequences in UniProtKB and UniProt Reference Proteomes as well as in the metagenomic MGnify database [19, 20].

### Clusters with sequence similarities to known adhesive domains

The Jackhmmer search using the putative adhesive region indicated 8 out of the 24 sequence clusters (cluster number 2, 6, 8, 15, 17, 19, 21, and 24) to be similar to known adhesive domain sequences (Table 4). These sequence similarities were confirmed by the DALI search with the AlphaFold predicted structure models (Figure 4a-i, S2a-i).

**Figure 4:**
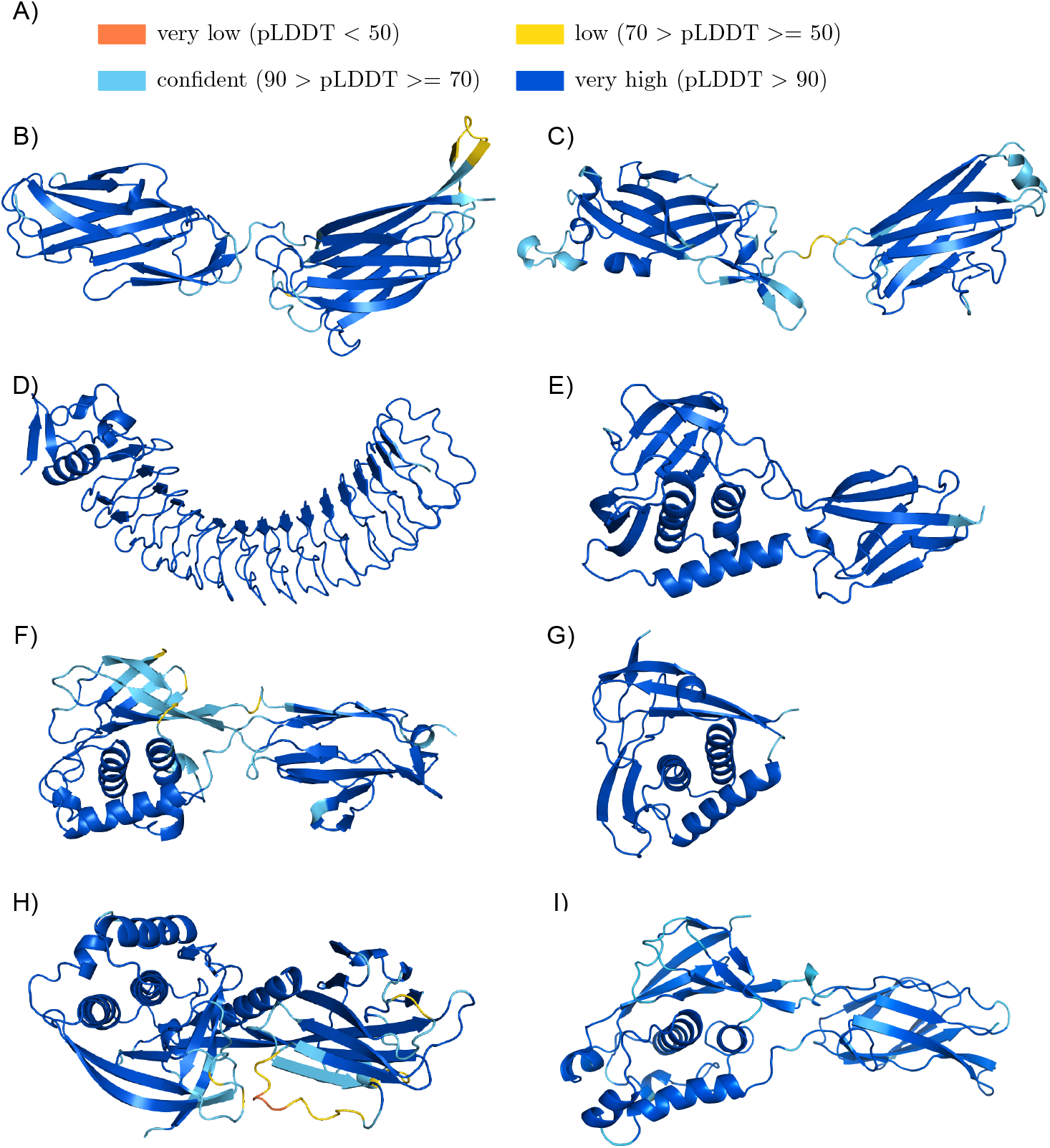
Structure models of cluster groups with sequence similar adhesive domains found by Jackhmmer: (a) Colour legend representing the quality of the AlphaFold models. Structure models of the potential adhesive domain of (b) cluster 2 (UniProtKB:K8EVB1/117-417), (c) cluster 6 (UniProtKB:V2XMF4/80-366), (d) cluster 15 (UniProtKB:A0A4U7JL97/37-420), (e) cluster 8 (UniProtKB:R3TX93/40-276), (f) cluster 17 (UniProtKB:A0A2Z4U801/240-508), (g) cluster 19 (UniProtKB:S0JHJ4/32-199), (h) cluster 21 (UniProtKB:A0A373LEL7/30-196) and (i) cluster 24 (UniProtKB:B1BZ86/36-324). The figures were produced using Pymol [26].

**Table 4:** Information about clusters with detectable sequence similarities to known Pfam adhesive domains. The similar Pfam adhesive domains found with Jackhmmer are indicated under ‘Domain overlap’. Other information are the homologous sequence hits per cluster in the UniProtKB, UniProt Reference Proteomes and MGnify database as well as the cluster size (‘Seqs #’) and the average protein prediction score per cluster (‘Avg. Score’).

| Nr. | Avg. Score | Seqs. # | Domain overlap | UniProtKB | UniProt Ref. Proteomes | MGnify |
| --- | --- | --- | --- | --- | --- | --- |
| 2 | 0.93 ± 0.06 | 8 | Big_8 (PF17961), Collagen_bind (PF05737) | 487* | 19* | 650* |
| 6 | 0.86 ± 0.05 | 6 | Collagen_bind (PF05737) | 276* | 66* | 1,174* |
| 8 | 0.84 ± 0.09 | 28 | TED (PF08341) | 1,149 | 222 | 48,395 |
| 15 | 0.78 ± 0.11 | 8 | LRR_4 (PF12799) | 45,022 | 2,793 | 170,274 |
| 17 | 0.78 ± 0.05 | 47 | TED (PF08341) | 1,170 | 170 | 54,096 |
| 19 | 0.77 ± 0.07 | 39 | TED (PF08341) | 954 | 148 | 28,283 |
| 21 | 0.74 ± 0.03 | 7 | TED (PF08341) | 56** | 16** | 3,617** |
| 24 | 0.71 ± 0.05 | 13 | TED (PF08341) | 1,223 | 214 | 53,312 |

\* C-terminal domain
\*\* N-terminal domain

#### Cluster 2

The predicted structure of cluster 2 representative sequence (UniProtKB:K8EVB1) shows that it contains two distinct domains (Figure 4b). For the N-terminal domain Jackhmmer identified a Big_8 domain (Pfam:PF17961), which is found in a variety of bacterial adhesins such as the *Staphylococcus aureus* proteins FnBPA, ClfA and ClfB. A DALI search with the N-terminal domain strengthens the Jackhmmer results as the best hit is the N-terminal Big_8 domain of the FnBPA binding region (PDB:4b60:B). For the C-terminal end of the C-terminal domain of cluster 2 the Jackhmmer search indicates an overlap to the Collagen_bind domain (Pfam:PF05737). The second domain structure yields with the DALI search the *Streptococcus gordonii* adhesin Sgo0707, where the structure superposes to the Sgo0707_N2 domain (Pfam:PF20623) (PDB:4igb:B). The subsequent DALI hits are the SdrG_C_C adhesive domain (Pfam:PF10425) (PDB:4jdz:A) (Figure S2b). The SdrG_C_C domain is also often found associated with the Big_8 domain in known adhesins and is likely to be homologous to the Collagen_bind domain. Cluster group 2 includes the *Enterococcus faecalis* protein with the gene name EF2505 or Fss2, which was described to bind to fibrinogen and to play an important role in the adherence and virulence of *E. faecalis* [21]. Thus we propose that proteins in cluster 2 are likely to bind fibrinogen or other animal extracellular matrix proteins.

#### Cluster 6

For cluster 6 AlphaFold predicted a structure composed of two domains (Figure 4c). The DALI search indicates, similar to cluster 2, for the N-terminal domain a Big_8 domain as the top hit (PDB:5cf3-A) and for the C-terminal domain a Collagen_bind domain as the best match (PDB:2z1p-A) (Figure S2c), thereby confirming the Jackhmmer search results. The Big_8 domain functions together with the Collagen_bind domain as supra-domain, enabling the binding to collagen by the collagen hug binding mechanism [22].

#### Cluster 15

Cluster group 15 includes the *Listeria monocytogenes* InternalinJ (UniProtKB: Q8Y3L4), for which the crystal structure of the adhesive domain exists (PDB:3bz5:A) [23], which was found using DALI with the predicted AlphaFold structure, reflecting the high accuracy of AlphaFold (Figure 4d, S2d). The adhesive domain is formed by a series of Leucine Rich Repeats that are not matched by the LRR_4 family (Pfam:PF12799) in Pfam. Proteins related to this class are one of the most prevalent that we found with over 170,000 homologues identified in the MGnify protein sequence database.

#### Cluster 8

The Jackhmmer search as well as the DALI search with the AlphaFold prediction indicate cluster 8 being a class II TED (Pfam:PF08341) adhesive domain (PDB:6fx6:A) (Figure 4e, S2e), whose binding partner is unknown [24, 25]. The TED adhesive domain is categorized into a class I and class II TED domain, depending on an additional N-terminal indel forming an alpha helix or an additional C-terminal indel folding into a beta-sandwich, respectively [24]. Cluster group 8 includes the fibrinogen binding *E. faecalis* Fss3 protein (UniProtKB:Q833P7) as well as an *E. faecalis* protein (UniProtKB:Q831Z5) encoded by the virulence associated EF2347 gene [21].

#### Cluster 17

The Jackhmmer search indicates cluster 17 to also be distantly related to the TED adhesive domain [25]. The DALI results with the best hit being a class II TED domain (PDB:6fwv) (Figure 4f, S2f) strengthens this hypothesis. To confirm that cluster 17 is a class II TED domain, we extended the representative sequence above the 400 residues limit by around 100 residues in order to include the characteristic C-terminal indel.

#### Cluster 19

Again, a TED domain was indicated by the Jackhmmer search for cluster 19, and supported by the DALI results, which indicated a class II TED domain (PDB:6fx6:A) (Figure 4g, S2g). Analysing the domain topology showed that there exists no seven-stranded beta-sandwich insertion, which is present in class II TED domains, whereby two beta-strand (A’ and B’) missing in class I TED domains are present [24]

#### Cluster 21

The predicted structure of cluster 21 showed two distinct domains (Figure 4h, S2h). The Jackhmmer search again indicated a TED domain. This is supported for the N-terminal domain by the DALI results, which indicated a class II TED domain. As in cluster 19, the seven-stranded beta-sandwich insertion is not present, but the two beta-strands A’ and B’. The C-terminal domain structure includes ten beta-strands and two longer alpha helices. The best DALI hit for the C-terminal domain is a stalk-like structure (PDB:3kpt:A), which only aligns to the beta-strands of the domain. Interestingly, the N-terminal TED-like domain is predicted to interact with the C-terminal domain.

#### Cluster24

The Jackhmmer search as well as the DALI results indicate a TED domain for cluster 24 (PDB:6fwy:B). The existing beta-stranded insertion clearly characterises this TED domain as class II TED domain (Figure 4i and S2i).

The sequences of five of the eight above described clusters show similarity to known TED adhesive domains. TED domains were previously described to show a high sequence diversity [25].

### Clusters with structure models indicating role in adhesion function

The cluster groups 1, 3, 4, 5, 10, 12 and 13 don’t yield persuasive Jackhmmer matches (Table 5), but their predicted structures resemble structures of known adhesive domains or adhesion associated domains, suggesting that these clusters might be novel domains with potential binding functions.

**Table 5:** Investigation of the potential novel adhesive domains: This table lists information regarding overlapping known Pfam domain families found with Jackhmmer (Domain overlap) and the abundance in the UniProtKB, UniProt Reference Proteomes and MGnify database for the N-terminal sequence clusters. The ‘Seqs #’ column represents the cluster size and the ‘Avg. Score’ column the average prediction score of the proteins per cluster.

| Nr. | Avg. Score | Seqs. # | Domain overlap | UniProtKB | UniProt Ref. Proteomes | MGnify |
| --- | --- | --- | --- | --- | --- | --- |
| 1 | 0.93 ± 0.09 | 7 |  | 256 | 38 | 569 |
| 3 | 0.91 ± 0.05 | 9 |  | 502 | 122 | 1,124 |
| 4 | 0.89 ± 0.06 | 8 |  | 145 | 32 | 157 |
| 5 | 0.87 ± 0.06 | 11 | MBG (PF17883)<br>(C-terminal) | 53 | 24 | 125 |
| 10 | 0.82 ± 0.07 | 10 |  | 2,359 | 551 | 6,668 |
| 12 | 0.82 ± 0.05 | 6 | Cthe_2159<br>(PF14262) | 495 | 65 | 1,495 |
| 13 | 0.8 ± 0.08 | 8 |  | 444 | 47 | 1,180 |

#### Clusters with jelly-roll resembling structure predictions

The structure predictions for cluster 4 and 5 are jelly-roll like structures (Figure 5a-c).

**Figure 5:**
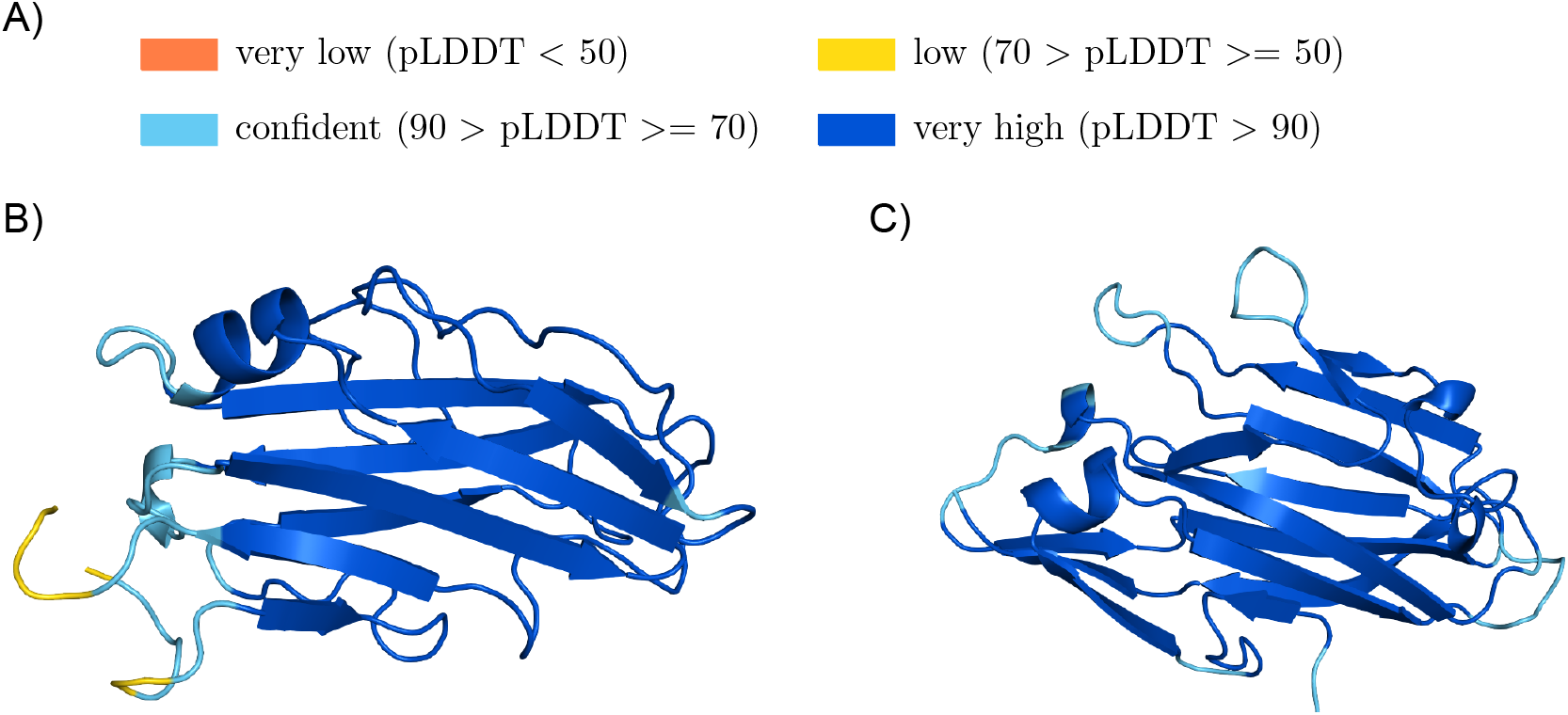
Potential novel adhesive domains with jelly-roll like predicted structures: (a) Colour legend representing the quality of the AlphaFold models. Structure models of the potential adhesive domain of (b) cluster 4 (UniProtKB:B0S3M8/153-316) and (c) cluster 5 (UniProtKB:A0A5R8Q9T8/68-240). The figures were produced using Pymol [26].

The best DALI hit for cluster group 4 was a GramPos_pilinBB domain of the RrgB pilus backbone (PDB:2×9x:A) (Figure 5b, S3b).

The Jackhmmer search for cluster group 5 showed an overlap with the MBG stalk domain at the C-terminus of the sequence, which was trimmed off, leaving a significant N-terminal sequence region for which the structure was predicted. The top DALI matches are related to surface adhesins with the best match being a *Clostridium perfringens* pilin protein (PDB:5xcb:A) with the GramPos_pilinBB domain (Pfam:PF16569) (Figure S3c).

The Evolutionary Classification Of protein Domains (ECOD) databases classifies the Gram-Pos_pilinBB domain under the topology named ‘Common fold of diphtheria toxin/transcription factors/cytochrome f’ [27]. This category also includes the adhesive domains SdrG_C_C and Collagen_bind. The Collagen_bind adhesive domain is a well studied jelly-roll structure [28], which is composed of two antiparallel beta-sheets and two short alpha-helices. Both AlphaFold structures for cluster 4 and 5 fold into jelly-roll like structures and show a high similarity to the Collagen_bind domain (PDB:1amx-A). The structure surfaces seem to provide a groove on the beta-sheets, indicating a potential collagen binding site [28]. However, the Collagen_bind domain alone has a 10-fold lower collagen binding affinity compared to the collagen hug binding mechanisms based on the Big_8:Collagen_bind supra-domain [22]. The similarity to the Collagen_bind structure, but also the N-terminal protein position distal to the cell surface anchor strongly suggests adhesive function for clusters 4 and 5.

#### Beta-solenoid fold cluster structure models

Clusters 1, 10 and 12 are predicted to fold into beta-solenoid structures (Figure 6a-d). The predicted structures for cluster group 1 as well as 12 are most similar to the binding region of the serine-rich repeat protein (SRRP) from *Lactobacillus reuteri* strain 100-23C (PDB:5ny0:A) (Figure S4b,c), being described to bind to epithelial cells, pectic acids and to play a role in the biofilm formation [29]. The Jackhmmer search for cluster 12 already indicated after the second iteration a distant relation to the carbohydrate binding Cthe_2159 (Pfam:PF14262) domain which is part of the Pectate Lyase superfamily, whereas the DALI search clearly indicated the highest similarity to the *L. reuteri* SRRP adhesive region [29]. The SRRP protein is not part of any existing Pfam family. Although we limited the clustering sequence to 400 residues we investigated whether the domains were longer with AlphaFold and extended it in the case of cluster 1 to about 800 residues. Interestingly, cluster 1 is found on a *Staphylococcus epidermidis* protein with a SasG_G5-E stalk (UniProtKB:A0A3G1RMM4), which was so far only found associated with the Bact_lectin adhesive domain (Pfam:PF18483) in *S. epidermidis* and *S. aureus* SasG homologues [30]. In our previous study we discussed the possibility that an adhesive domain can function with any arbitrary stalk [7]. The described example underlines this hypothesis and furthermore indicates a possible transfer of the adhesive domain onto a given stalk increasing the adhesins variability. The SasG_G5-E stalk is also described to promote biofilm formation [30].

**Figure 6:**
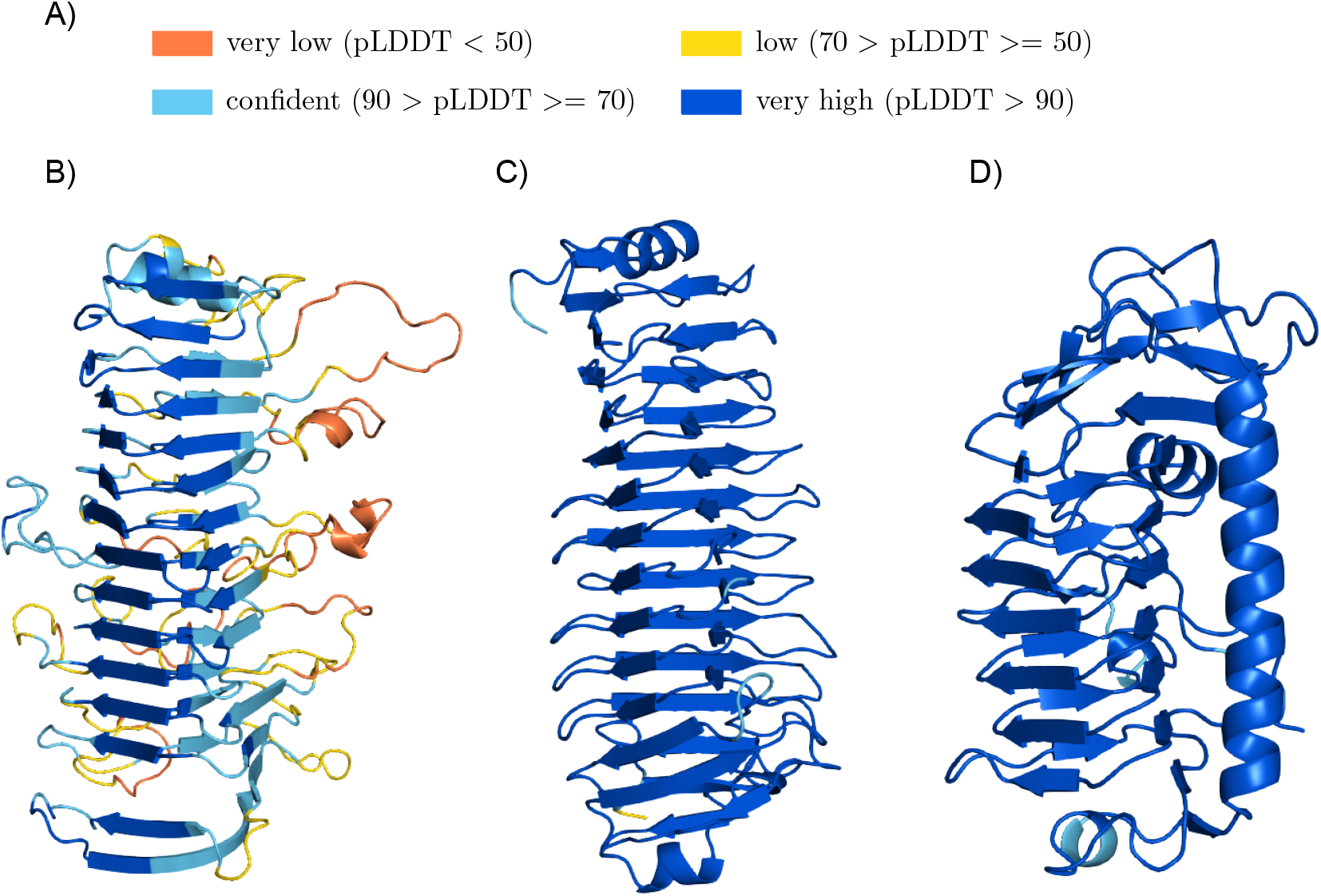
Predicted structures for clusters with potential novel adhesive domains, whose structure, but not sequence, seem to be related to known adhesive domains: (a) Colour legend representing the quality of the AlphaFold models. Structure models of the potential adhesive domain of (b) cluster 1 (UniProtKB:A0A2Z6T9E9/185-718), (c) cluster 12 (UniProtKB:A0A099WCN8/48-417) and (d) cluster 10 (UniProtKB:A0A0F7RLJ7/46-326). The figures were produced using Pymol [26].

Cluster group 10 resembles an ice binding domain (PDB:4nuh:A) (Figure S4d), where the representative protein sequence (UniProtKB:A0A0F7RLJ7) is identical to a *Bacillus anthracis* protein (UniProtKB:A0A384LNE7) with the gene name BA_0871 or BASH2_04951, which was described to be collagen binding and to be linked to the bacterial pathogenicity [31].

#### Remaining clusters with structure models indicating potential adhesion function

The best DALI hit for the cluster 3 structure model (Figure 7b) is an N-terminal helical domain of a group B streptococcus immunogenic bacterial adhesin named BibA (PDB:6poo:A) (Figure S5b), which superposes with the N-terminal alpha helices of the structure model [32]. Running the DALI search for the domain in the middle of the structure model separately, results in the *S. aureus* SdrD adhesive protein (PDB:4jdz:A), where the structure model superposes with the SdrD_B stalk domain (Pfam:PF17210). The cluster 3 includes one *Streptococcus merionis* protein (UniProtKB:A0A239SMH4), which is encoded by the bca gene. The bca gene has been shown to be involved in the initial stage of Group B Streptococcus infection [33], suggesting adhesion function.

**Figure 7:**
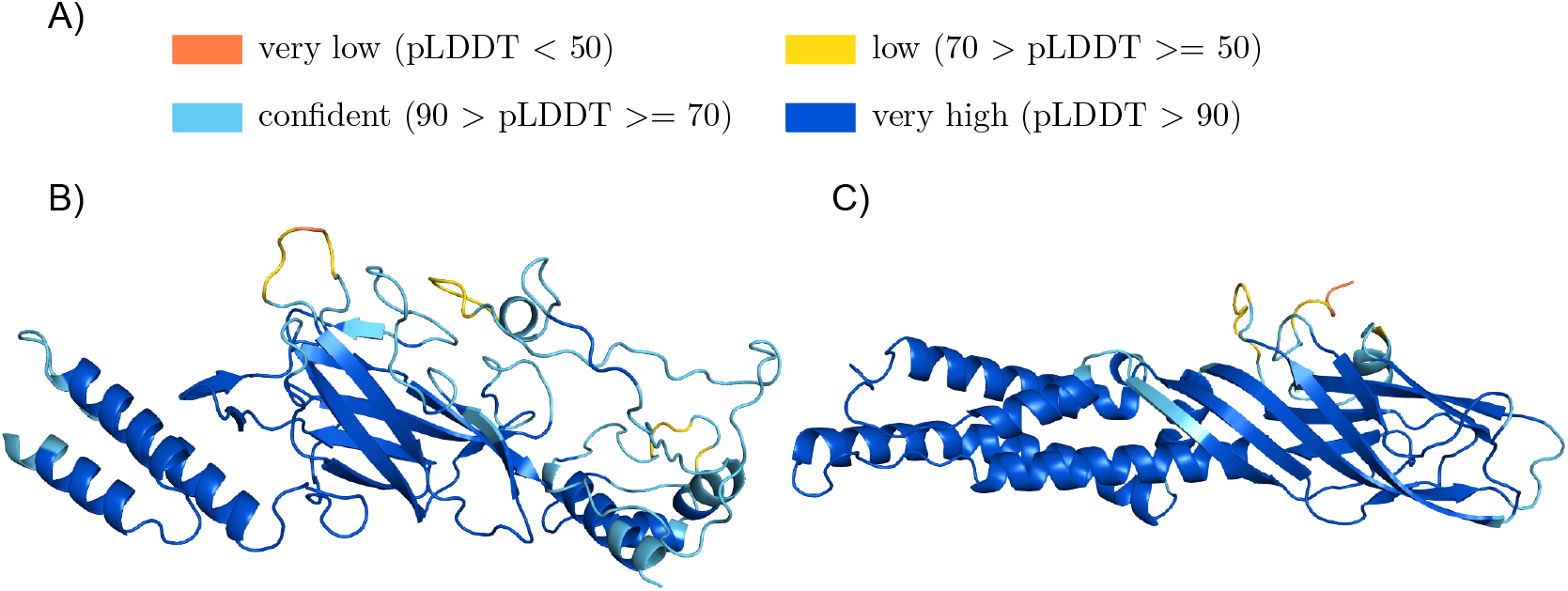
Clusters with AlphaFold structures showing ambiguous adhesion function: (a) Colour legend representing the quality of the AlphaFold models. Structure models for (b) cluster 3 (UniProtKB:A0A1Q8E8C7/76-420) and (c) cluster 13 (UniProtKB:A0A069CUH0/64-357). The figures were produced using Pymol [26].

The best DALI match for cluster 13 is the human integrin alpha-5 protein (PDB:7nxd:A), followed by the best bacterial match being the N-termini of the *S. gordonii* adhesin Sgo0707 (PDB:4igb:B) (Figure 7b, S5c). Here, the structure model aligns to the Sgo0707_N2 domain. The cluster includes a *E. faecalis* protein (UniProtKB:Q82YW8) encoded by the EF3314 gene, which was described to contribute to the virulence properties of this pathogen [34].

We created new putative adhesive Pfam domain families for the clusters 1, 4, 5, 10, 12, 13 and 24. Clusters 1 and 12 were combined into a single cluster. The Pfam identifiers can be found in supplementary table S4. The common domain architectures of proteins with these potential novel members of adhesive domain families are shown in figure 8.

**Figure 8:**
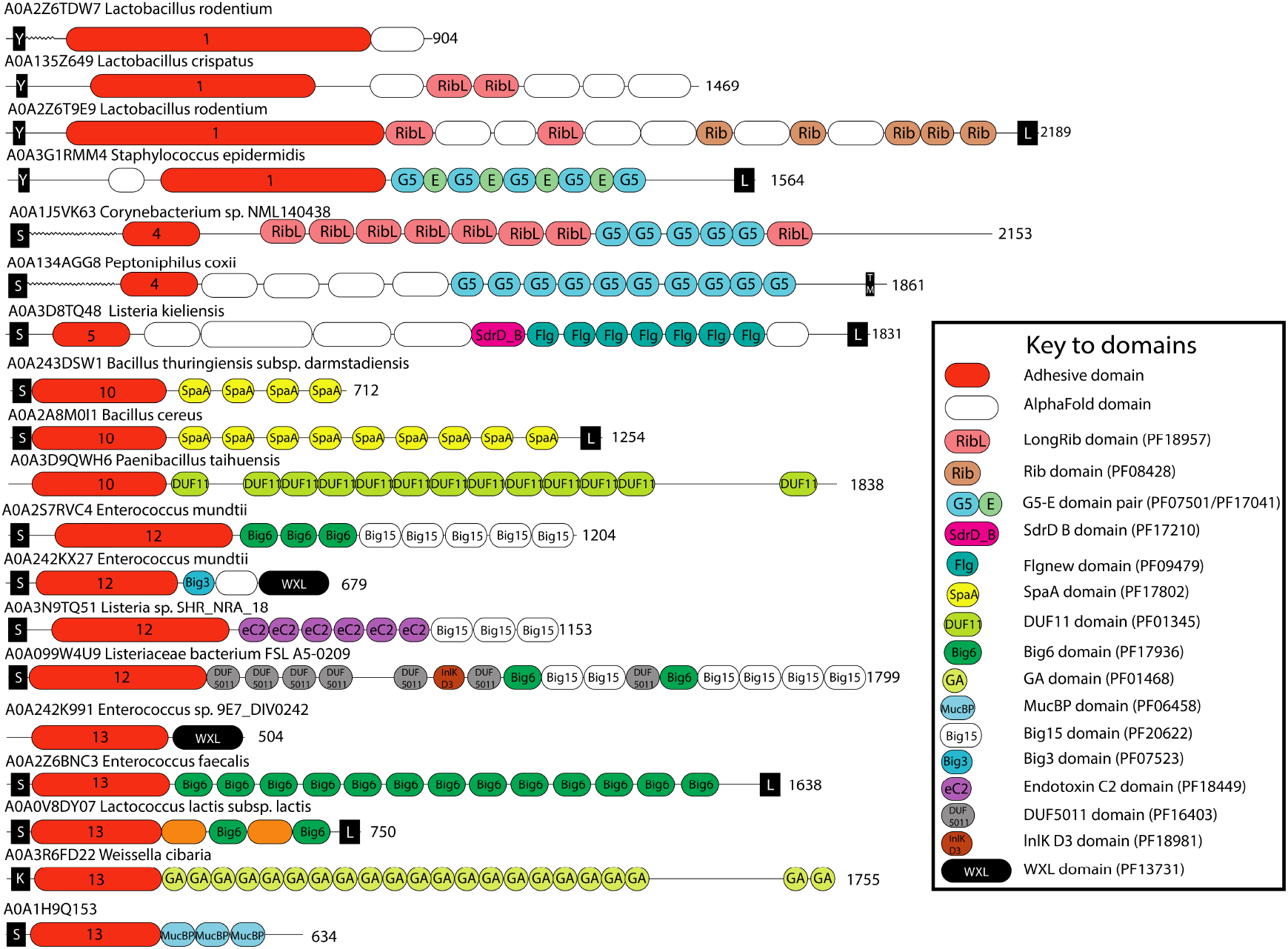
Examples of protein architectures in which the potential novel adhesive domains can be found: The potential novel adhesive domains are annotated in red, labelled with the cluster number. The white domains (‘AlphaFold domains’) are domains found in the AlphaFold structure model of each protein, which do not correspond to existing Pfam domain families.

### Clusters with stalk-like domains

The Jackhmmer search for clusters 11, 14, 20 and 22 indicated known stalk domains (Table 6). The structure predictions support these results (Figure 9 and figure S6), suggesting the N-terminal region to be composed of stalk domains without a functional N-terminal adhesive domain. As discussed in our previous work, the boundary between adhesive and stalk domains is not always clear, opening the question whether stalk domains can develop binding functions [7]. Additionally, we can find stalk domain structures with similarities to adhesive domains, for example the DUF11 domain family (Pfam:PF01345) has similarities to the Collagen_bind adhesive domain structure (Pfam:PF05737).

**Figure 9:**
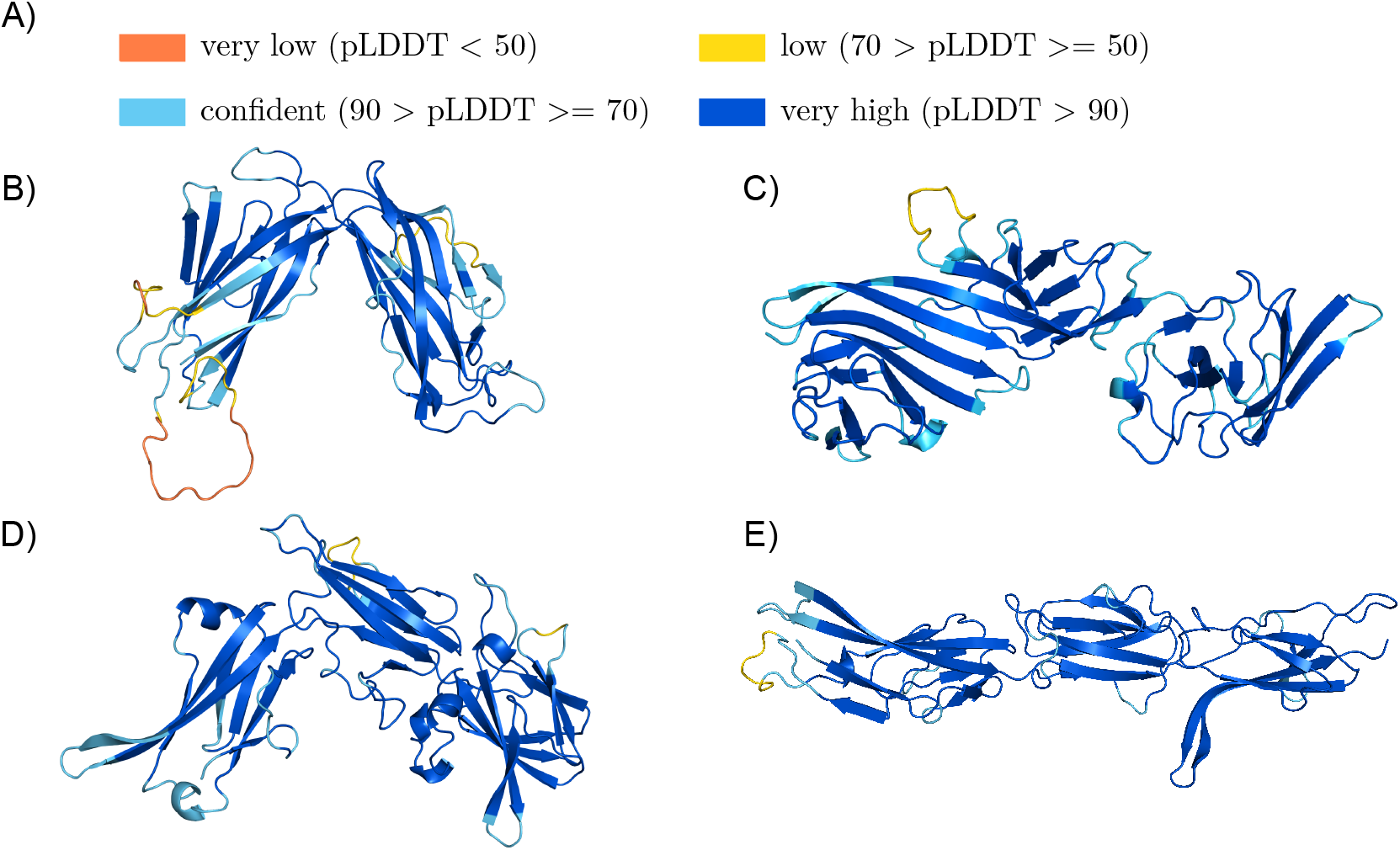
Cluster with AlphaFold structure models with stalk characteristics: (a) Confidence of AlphaFold models colour legend. Structure models for b) cluster 11 (UniProtKB:A0A2V5K856/38-347), c) cluster 14 (UniProtKB:A8MK03/34-337), d) cluster 20 (UniProtKB:A0A1I6IKX0/39-366) and e) cluster 22 (UniProtKB:R7HBU9/38-352).

**Table 6:** Information about stalk-like domain clusters: The table contains the cluster number, average prediction score and number of sequences per cluster (‘Seqs #’). The Pfam stalk domains found with Jackhmmer are indicated under ‘Domain overlap’. Information is also included about the number homologous sequence hits per cluster in the UniProtKB, UniProt Reference Proteomes and MGnify database.

| Nr. | Avg. Score | Seqs. # | Domain overlap | UniProtKB | UniProt Ref. Proteomes | MGnify |
| --- | --- | --- | --- | --- | --- | --- |
| 11 | 0.82 ± 0.05 | 10 | DUF11 (PF01345) | 476* | 87* | 1,237* |
| 14 | 0.8 ± 0.04 | 22 | TIG (PF01833) | 988* | 207* | 2,190* |
| 20 | 0.75 ± 0.03 | 10 | Big_2 (PF02368) | 915* | 228* | 9,914* |
| 22 | 0.73 ± 0.02 | 6 | Big_2 (PF02368) | 109* | 35* | 3,285* |

\* N-terminal domain

A second possible function of these proteins is to act as steric regulators altering the access of other adhesive proteins to binding partners. One example is the *S. aureus* periscope protein SasG, which is suggested to block the binding of proteins located closer to the cell surface from interacting with host cell fibrinogen [9].

The Jackhmmer search for cluster 11 resulted in the known DUF11 (Pfam:PF01345) stalk domains. The predicted structure shows two distinct domains (Figure 9b). The DALI search indicated for the N-terminal domain a similarity to the stalk-like structure of an Integrin alpha-X protein (PDB:4nen:A) (Figure S6b) and for the C-terminal domain a similarity to the BcpA major pilin subunit (PDB:3rkp:A).

The sequence as well as the structure resemble a TIG stalk domain for the C-terminal domain of cluster 14, indicated by Jackhmmer and the DALI search (PDB:5l5g:D) (Figure 9c, S6c). The N-terminal part is composed out of two subdomains, which seem to mirror each other. This symmetry could be based on an internal duplication event. The best DALI hit with a Z-score of 8.1 for the N-terminal part was a plexin-C1 protein (PDB: 6vxk:D), where the cluster aligns to the two TIG domains in the protein, whereby only the N-terminal subdomain superposes well (Figure S6C). But separately, both subdomains of the N-terminal part superpose reliably to a TIG domain, suggesting the N-terminal part to be related to a combination out of two TIG domains, which might have developed further. A groove on the surface of the structure model indicates that the N-terminal part might have developed a binding function (Figure 9c).

The structure model for cluster 20 presents three domains (Figure 9d). The Jackhmmer search already indicated a Big_2 stalk domain, which is supported for the N-terminal domain within the top DALI results (PDB:2l04-A) (Figure S6d). The middle domain resembles a stalk-like structure in Intimin_C (PDB:1f00:I) and the best DALI hit for the C-terminal domain was the CfA/I fimbrial subunit A (PDB:6k73:B).

The sequence of cluster 22 also indicated a Big_2 stalk domain. The predicted structure again represents three domains, of which the N-terminal and C-terminal domain are most similar to the stalk domains PKD_4 (PDB:4u7k:G) and Big_2 (PDB:4hu8-C) respectively (Figure 9e, S6e). The best DALI hit for the domain in the middle is a monooxygenase (PDB:1yew:E) closely followed by the I-set immunoglobulin-like stalk domain (PDB:5aea:A).

### Clusters with ambiguous function

The Jackhmmer and DALI search could not indicate explicit binding function for cluster group 7, 9, 16, 18 and 23 (Table 7, Suppl. figure S7).

**Table 7:**
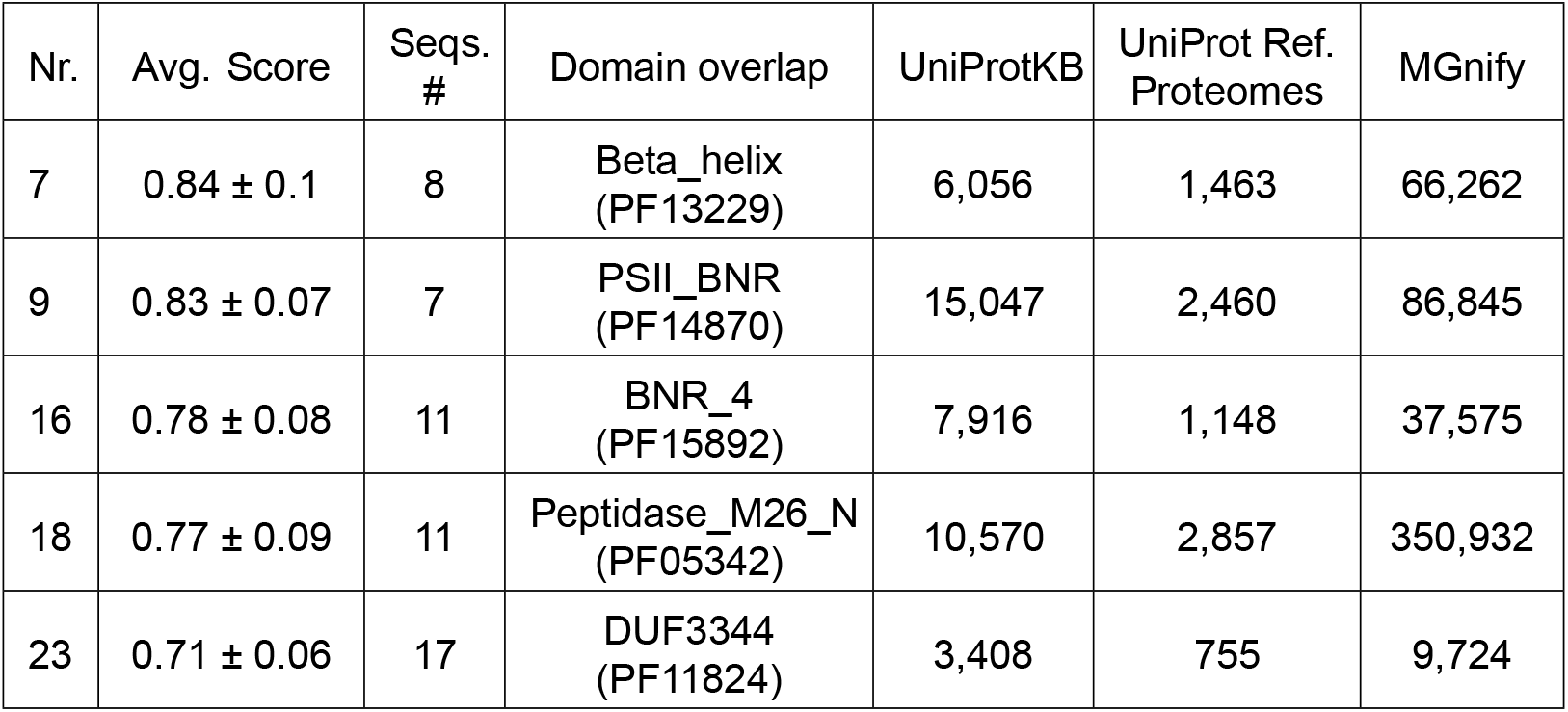
Information about clusters of ambiguous function: Information regarding overlapping known Pfam domain families found with Jackhmmer(Domain overlap) and the abundance in the UniProtKB, UniProt Reference Proteomes and MGnify database are listed for the N-terminal sequence clusters. The ‘Seqs #’ column represents the cluster size and the ‘Avg. Score’ column the average prediction score of the proteins per cluster.

| Nr. | Avg. Score | Seqs. # | Domain overlap | UniProtKB | UniProt Ref. Proteomes | MGnify |
| --- | --- | --- | --- | --- | --- | --- |
| 7 | $0.84 \pm 0.1$ | 8 | Beta_helix (PF13229) | 6,056 | 1,463 | 66,262 |
| 9 | $0.83 \pm 0.07$ | 7 | PSII_BNR (PF14870) | 15,047 | 2,460 | 86,845 |
| 16 | $0.78 \pm 0.08$ | 11 | BNR_4 (PF15892) | 7,916 | 1,148 | 37,575 |
| 18 | $0.77 \pm 0.09$ | 11 | Peptidase_M26_N (PF05342) | 10,570 | 2,857 | 350,932 |
| 23 | $0.71 \pm 0.06$ | 17 | DUF3344 (PF11824) | 3,408 | 755 | 9,724 |

The Jackhmmer search indicated a Beta_helix (Pfam:PF13229) for cluster 7 and a Peptidase_ M26_N (Pfam:PF05342) domain for cluster 18. The structure models of both domains resemble carbohydrate binding pectate lyase adhesive domains. But the top DALI hit for cluster 7 is a lacto-N-biosidase (PDB:6kqs:A) and for cluster 18 a putative immunoglobulin protease (PDB:3n6z:A), suggesting potential catalytic functions [35].

Jackhmmer indicated a BNR_4 (Pfam:PF15892) domain for cluster 16 and the top DALI hit for this cluster is a human integrin alpha-IIb protein, which binds among others to fibrinogen. The InterPro database has a Fucose_binding_lectin domain annotated (PDB:1iub), suggesting cluster 16 to have adhesion function [36, 37]. The structure model of cluster 9 resembles cluster 16, whereby the top DALI hit is a virginiamycin B lyase (PFB:2z2o:B), again suggesting catalytic function [38].

The best DALI hit for cluster 23 was the P_proprotein domain (Pfam:PF01483) of a protease, which is common to be located downstream of a catalytic domain (PDB:3hjr:A) [39].

The results of the above described clusters suggest the clusters to play a catalytic role, not ruling out binding abilities. Given that catalytic domains often also have adhesive function to bind to their substrate, it is challenging to differentiate between catalytic and adhesive domains and also between fibrillar adhesins and as we call them ’fibrillar enzymes’. Fibrillar enzymes are also composed out of repeating domains, but have an enzymatic related domain instead of an explicit adhesive domain (Figure 10). Nevertheless, the enzymatic region can be able to have binding functions, as described above.

**Figure 10:**
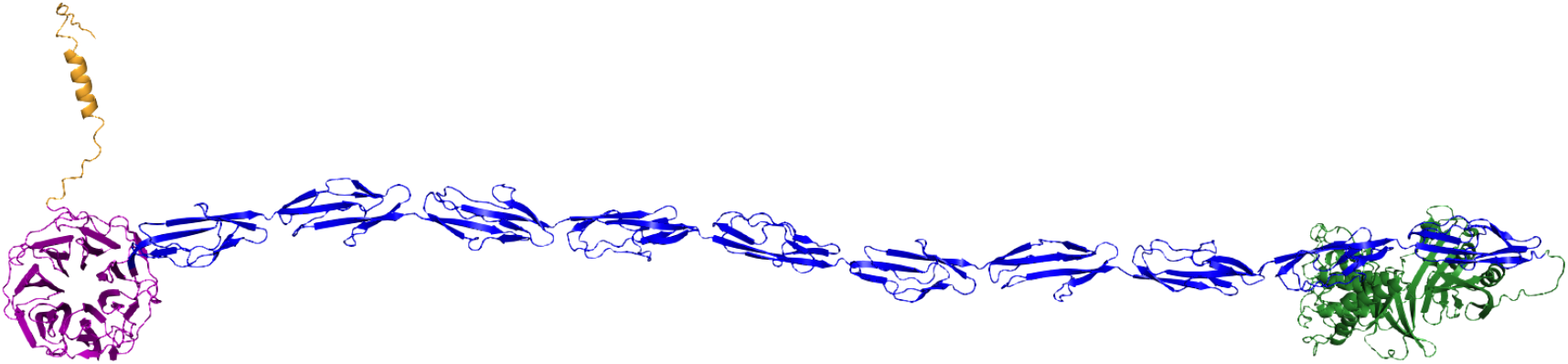
Example of a potential fibrillar enzyme: Predicted structure of a potential fibrillar enzyme (UniProtKB:C6CUY3). The structure was predicted using the AlphaFold colab notebook, where three sequence chunks (residues 1-600, 401-1000 and 901-1500) were predicted separately with overlapping regions, which were combined using pymol. The potential sorting signal region is coloured in yellow, the potential catalytic domain in violet, the Flg_new stalk domains in blue and the anchor region in green. This protein belongs to the sequence cluster 9. This figure were produced using Pymol [26].

## Discussion

Novel pathogens are emerging constantly with uncharacterized host cell interaction mechanisms. Homologous virulence associated proteins with known adhesive domains are the first step towards an understanding of the pathogenicity of these bacteria. But adhesive domains evolve quickly and are highly variable. Hence, detection mechanisms independent from known adhesive domains are important. In this study we developed a random forest based discovery approach to detect FA-like proteins. We applied the approach on the Firmicute and Actinobacteria UniProt Reference Proteomes, yielding over 6,500 confidently predicted FA-like proteins. With the characterization of FA-like proteins we could identify a variety of notable features which we could use for machine learning. The known stalk and adhesive domains are the strongest feature in the classification decision approach. This bias is due to the positive training data, which was selected from the prior domain-based discovery approach [7]. Other strong features were the protein length, which is required to overcome the bacterial cell surface, and the amino acid composition of the protein sequences. Here, particularly threonine was strongly over-represented in the positive training data set compared to the negative training data set, raising the question about what role it plays in bacteria-host interactions. One explanation could be that despite the most commonly phosphorylated amino acids being histidine and aspartate, serine/threonine/tyrosine phosphorylation in bacteria was shown [40, 41]. Additionally, phosphorylation during the adherence step was described, connecting it to the regulation of the bacterial virulence [41, 42]. Fibrillar adhesins are cell surface proteins and so we selected the existence of a cell wall anchor motif or domain as an additional feature. Although an anchor was found in only around half of the proteins of the positive training data. One reason could be that there are many unknown anchor motifs or domains, which still need to be investigated, or non-classical secretion mechanisms [43]. We also found many examples of potential fibrillar adhesins where the stalk region ranges to the C-terminus. These proteins might be able to interact with other cell surface proteins in order to be projected away from the cell wall. Implementing the selected identification properties in the random forest classification approach and applying it to the Firmicute and Actinobacteria UniProt Reference Proteomes led to over 6,500 confidently detected FA-like proteins. This indicates that more than 5,000 of them were missed by the domain based discovery approach detected in our previous study [7]. More importantly, with our new machine learning discovery approach FA-like proteins are predicted that lack known adhesive or stalk domains enabling us to discover novel protein domain families.

To verify the random forest prediction approach we further studied the predicted FA-like proteins lacking a known adhesive domain, but with known stalk domains. When investigating the sequence clusters representing an annotation gap N-terminal to known stalk domains, similar Pfam domain sequence matches could be found for many of the described clusters using Jackhmmer. This suggests that the Pfam domain families could be expanded to include these sequences or novel domain families related to the overlapping domain families can be created. We have taken advantage of the recent release of the AlphaFold2 software to validate our machine learning approach as well as use it to refine predictions of adhesive domains in our predicted fibrillar adhesins. Given that the predicted structures confirm the Jackhmmer results, highlights the high accuracy of the structure prediction method AlphaFold2. We see many new opportunities to use large scale structure predictions to identify and investigate the components of the bacterial cell surface that are likely to interact with the host.

While further investigating the described N-terminal sequence clusters the difficulty to differentiate between fibrillar adhesins and the newly discovered class of fibrillar enzymes was shown. Given that fibrillar enzymes can play an important role in the bacterial pathogenesis as well, they have comparable characteristics to fibrillar adhesins in terms of being long surface proteins with a stalk and that several of the enzymes can have binding functions. This impedes the differentiation of these two protein classes by our identification features and so far we have not included any property to differentiate between adhesive and enzymatic domains. Nevertheless, the prediction score and the cluster size can together give an assessment about the reliability. The analysed clusters with the higher sequence number or higher prediction score are mostly potential adhesive domains.

Except for cluster 3 and 13, the sequences or predicted structures of the other potential adhesive domain clusters are similar to known adhesive domains. Additionally, the potential novel adhesive domains as well as the clusters related to known adhesive domains found with Jackhmmer verify the random forest based discovery approach. The high number of homologous sequences of these domains in the metagenomic MGnify database and in known pathogenic genera in the UniProt database underline their relevance. For the potential novel members of adhesive domain families discovered in the course of this study the predicted structure models and the DALI search results give a first understanding of their function and potential binding partners. AlphaFold2 and AlphaFold-Multimer opens up further ways to predict the structures of fibrillar adhesins-target protein complexes [44]. We believe that we are at the beginning of a new age of discovery where computational analyses will lead to fundamental improvements in our understanding of microbial host interactions.

## Materials and Methods

### Training data selection

We selected as positive training data the FA-like proteins of Actinobacteria and Firmicutes discovered with the domain-based detection approach in our previous study [7]. Additionally, we included 25 additional FA-like proteins, which don’t have either a known adhesive and stalk domain, which were found in the literature or manually investigated. As negative training data we selected randomly non FA-like proteins in reference proteomes, in which FA-like proteins could be detected with the domain-based discovery approach. These are from the following nine organisms: *Bifidiobacterium subtile, Olsenella sp. oral, Slackia exigua, Streptomyces coelicolor, Staphylococcus aureus, Lactococcus lactis, Streptococcus gordonii, Listeria monocytogenes* and *Enterococcus faecalis.* The training set consists of a total of 3,332 proteins, of which half belong to the positive and the other half to the negative training data set. The training data can be found in the GitHub repository (see below).

### Identification features calculation and random forests classification

To search in the protein sequences for known adhesive, stalk and anchor domains, the collection of Pfam domain HMMs from our previous study was used [7]. Additional, the adhesive domain GspA_SrpA_N (Pfam:PF20164) and the stalk domains aRib (Pfam:PF18938), RibLong (Pfam:PF18957), SasG_E (Pfam:PF17041), GA-like (Pfam:PF17573), YDG (Pfam:PF18657), Lipoprotein_17 (Pfam:PF04200) and IgG_binding_B (Pfam:PF01378) were used. These HMMs were run against the protein sequences using the HMMER tool (version 3.1b2) with the gathering (GA) threshold option. Using regular expression, we searched within the C-terminal 50 residues of the protein sequences for the following sortase anchor motifs: ‘LPxTG’, ‘LPxTA’, ‘LPxTN’, ‘LPxTD’, ‘LPxGA’, ‘LAxTG’, ‘IPxTG’, ‘NPxTG’, ‘NPQTM’ (‘x’ can be any amino acid).

To identify highly similar tandem sequence repeats that may represent potential unknown stalk domains we applied the T-REKS software on the sequences using as parameters a minimum of 70% sequence identity, 50 residues as minimum length of the repeat region and 5 residues as minimum seed length [12].

Disordered regions were predicted using IUPred (IUPred2a) with the IUPred2 type ‘long’ for predicting long disordered regions [13]. Each residue with an IUPred score above 0.5 was counted as predicted disordered. The predicted disordered fraction was calculated using the percentage of predicted disordered residues from the total protein length.

The proportion of charged or hydrophobic amino acids per protein sequence was calculated using as charged amino acids glutamic acid (E), aspartic acid (D), lysine (K), arginine (R) and as hydrophobic amino acids alanine (A), isoleucine (I), leucine (L), methionine (M), phenylalanine (F), tryptophan (W), tyrosine (Y), valine (V).

The residue length was counted per the complete UniProt protein sequence.

The proportion for each amino acid per protein sequence was calculated and to evaluate the amino acid composition bias the relative entropy (Kullback-Leibler (KL) divergence) was calculated per protein sequence (S). Here, we quantify the difference between the observed frequency (P) per amino acid (i) compared to equally frequent amino acids, being 0.05 for 20 amino acids.

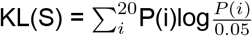

We calculated the identification features for the protein sequences of the training data. With the calculated feature data we trained a random forest classifier from sklearn ensemble methods with 50 trees with maximum 3 features per tree and random state 2 [45]. The Random Forest method takes the 30 features as input and outputs a score per protein between 0 and 1 with FA-like proteins scoring closer to 1.

The reliability curve was calculated for the applied random forest model on the training data set using calibration_curve from the sklearn calibration module and a 10-fold cross-validation approach.

For calculating the precision and recall of the model and generating the precision-recall curve, we generated a testing data set of 258 proteins, composed of 128 FA-like proteins and 130 non FA-like proteins. We artificially adapt the features of the testing set to have no adhesive or stalk domains, whereby all other features are retained. For these calculations the proteins of the testing set were excluded from the training data set. The precision and recall of the model as well as the precision-recall curve was calculated using the macro-average method to determine how the random forest model performs overall across the two classes: FA-like, non FA-like proteins. The precision-recall curve was also calculated using a cross-validation approach with the training set.

To use the random forest discovery approach, we provide the code in our GitHub repository (see below).

### FA-like proteins prediction for Firmicute and Actinobacteria UniProt Reference Proteomes

To apply our machine learning method against known Firmicute and Actinobacterial proteins we first gathered available sequences. The UniProt proteome identifier for all Firmicute and Actinobacteria Reference Proteomes were searched for in the UniProt website (release 2020_04). We collected the relevant sequences for these identifiers by searching in the knowledgebase under the bacterial reference proteomes (release 2020_03) for the identifier.

As described in the subsection ‘Identification features calculation and random forest classification’ we calculated the identification features for the Firmicute and Actinobacteria reference protein sequences and applied the trained random forest classification approach to score each protein.

### Analysing predicted FA-like proteins

We further analysed the predicted FA-like proteins by differentiating the prediction scores. The subcellular localizations of the predicted FA-like proteins were predicted using singularity (version 3.5.3) to run the PSORTb (version 3) Docker image (psortb_commandline_1.0.2.sif).

To find distantly related adhesive domains, a profile HMM-search with the known adhesive domains were conducted using an E-value threshold of 1.0.

Using profile HMM-search (version 3.1b2) with the gathering (GA) threshold option the Pfam database (version 33.1) was run against the sequences of the predicted FA-like proteins.

### Selecting potential functional sequences

To verify the machine learning approach, we were particularly interested in the predicted proteins with an annotation gap at the N-terminus, which might contain a missing functional domain. We focused on the N-terminus, because we showed in our previous study that the adhesive domain in FA-like proteins in Firmicutes and Actinobacteria is mostly found at the N-terminus [7]. We selected proteins with at least four known stalk domains, which lack a known adhesive domain and with no Pfam domain annotations within the first 20% of the protein length. Before finding homologous sequence groups, we deleted the selected protein’s first 20 residues to avoid clustering based on a potential signal peptide. We cut these sequences N-terminal to the first domain annotation, but no longer than 400 residues in order to try to avoid clustering based on potential stalk domains. We clustered those excised sequences into homologous sequence cluster groups using blastp all against all with an E-value threshold of 0.001, requiring a coverage threshold of 85% and an identity threshold of 25% [16]. For each cluster we calculated the reliability by averaging the random forest prediction scores of the proteins per cluster. We sorted the resulting clusters by average prediction score as well as sequences per cluster. We further investigated the 24 cluster groups with at least 5 homologous sequences.

To investigate the potential function of these sequence clusters, we chose one representative protein per cluster. To do so we aligned the N-terminal sequences per cluster and manually selected one representative sequence per cluster, which was used for the following investigations. For each representative sequence we searched the whole UniProtKB with Jackhmmer using the HMMER website (https://www.ebi.ac.uk/Tools/hmmer/search/jackhmmer) to find domain families related to the sequence clusters [17]. For cluster 5 we found a distant related stalk domain overlapping with the C-terminus of the representatives sequences, we trimmed off the sequence with the domain annotation and continued with the N-terminal sequence.

The structure for each representative sequence was predicted with AlphaFold2 using the Google colab repository provided by DeepMind (https://colab.research.google.com/github/deepmind/alphafold/blob/main/notebooks/AlphaFold.ipynb) [11]. Based on the predicted structure we selected the domain boundaries and cut the structure as well as sequence of each cluster accordingly (Suppl. Table S2). In most cases, we cut off disordered regions. In single cases, for cluster 4, we optimized the structure by cutting off a stalk domain-like C-terminal to focus on the potential adhesive domain and rerun the AlphaFold structure prediction. For cluster 1 and 17, we extended the sequence to include the whole domain.

To assess the quality of the models, AlphaFold stores the pLDDT confidence in the B-factor field of the output PDB files, which were used to colour the structure models by quality using Pymol [26]. To find out more about the function of the clusters, we searched with the predicted structure models, optimizised to the domain boundaries, for similar structures in the PDB database using DALI [46].

We created a HMM from the sequences per cluster based on the detected domains using hmm-build [17]. With these HMMs we searched against the metagenomic MGnify (release 2019_05), UniProt Reference Proteomes and UniProtKB (release 2021_01) databases for homologous sequences using an domain E-value threshold of 0.01 [19, 20]. From the UniProt website the Retrieve ID/Mapping tool was used to collect the organisms information to the UniProtKB matches.

## Data availability

The AlphaFold structure model as well as the Random Forest prediction results for the Firmicutes and Actinobacteria Reference Proteomes can be found in an institutional repository of the University of Cambridge (https://doi.org/10.17863/CAM.82322) [47]. We provide a GitHub repository (https://github.com/VivianMonzon/FAL_prediction), which includes the training data set and the code to run the Random Forest based FA-like protein prediction on a sequence of interest as well as a colab notebook (https://colab.research.google.com/github/VivianMonzon/FAL_prediction/blob/main/Colab/ML_FA_prediction.ipynb).

## Acknowledgment

We thank Dr. Aleix Lafita for his advice and his great knowledge on bacterial stalk domains.

