## Supplementary Material for "Large Scale Discovery of Microbial Fibrillar Adhesins and Identification of Novel Members of Adhesive Domain Families"

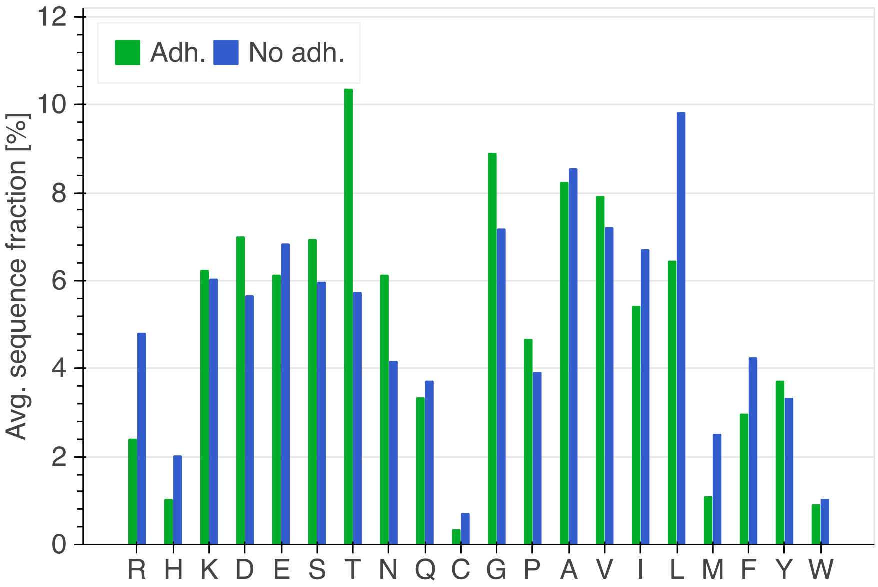


**Figure S1: Average amino acid fraction:** This bar plot shows for each amino acid the average fraction calculated per positive (Adh.) and negative (No adh.) training data set.

**Table S1: FAL per proteome counts:** These tables display the information to the reference proteomes with the highest fraction of predicted FA-like proteins with a score of 0.7 or above per reference proteome size for a) Firmicutes and b) Actinobacteria.

**a) Firmicutes**

| **Organism** | **Proteome ID** | **# FA-like proteins** | **Proteome size** | **Percentage** |
|---|---|---|---|---|
| *Listeria booriae* | UP000029844 | 57 | 3104 | 1.84 |
| *Culicoidibacter larvae* | UP000306912 | 43 | 2363 | 1.82 |
| *Granulicatella elegans* ATCC 700633 | UP000002939 | 20 | 1562 | 1.28 |
| *Weissella cryptocerci* | UP000292886 | 33 | 2588 | 1.28 |
| *Listeria monocyto-genes* serovar 1/2a | UP000000817 | 36 | 2844 | 1.27 |
| *Veillonella* sp. DORA_B_18_19_23 | UP000053161 | 11 | 870 | 1.26 |
| *Lactobacillus roden-tium* | UP000257317 | 18 | 1434 | 1.26 |

| | | | | |
|---|---|---|---|---|
| *Granulicatella balaenopterae* | UP000198556 | 24 | 1916 | 1.25 |
| *Clostridium* sp. CAG:433 | UP000018401 | 16 | 1308 | 1.22 |
| *Lactobacillus* sp. LL6 | UP000319179 | 22 | 1826 | 1.20 |

**b) Actinobacteria**

| **Organism** | **Proteome ID** | **# FA-like proteins** | **Proteome size** | **Percentage** |
|---|---|---|---|---|
| *Alloscardovia macacae* | UP000243657 | 18 | 1550 | 1.16 |
| *Rarobacter faecitabidus* | UP000315389 | 25 | 2325 | 1.08 |
| *Pseudoscardovia suis* | UP000216454 | 18 | 1732 | 1.04 |
| *Rarobacter incanus* | UP000316181 | 20 | 2018 | 0.99 |
| *Corynebacterium kutscheri* | UP000033457 | 20 | 2047 | 0.98 |
| *Micrococcales* bacterium KH10 | UP000198369 | 22 | 2348 | 0.94 |
| *Bifidobacterium primatium* | UP000229095 | 19 | 2029 | 0.94 |
| *Bifidobacterium indicum* LMG 11587 | UP000028569 | 12 | 1350 | 0.89 |
| *Atopobium vaginae* DSM 15829 | UP000005947 | 11 | 1243 | 0.88 |
| *Bifidobacterium bifidum* BGN4 | UP000006173 | 16 | 1834 | 0.87 |

**Table S2: Representative sequences:** this table lists the representative sequences and the regarding domain positions based on the AlphaFold structure prediction for the N-terminal annotation gaps of predicted FA-like proteins with minimum 4 stalk domains, lacking a known adhesive domain. The DALI search was conducted with the sequence regions listed here.

| Nr. | UniProt identifier | Domain info | Start position | End position |
|---|---|---|---|---|
| 1 | A0A2Z6T9E9 | | 185 | 718 |
| 2 | K8EVB1 | N-term | 117 | 256 |
| | | C-term | 257 | 417 |
| 3 | A0A1Q8E8C7 | | 76 | 420 |

| | | | | |
|---|---|---|---|---|
| 4 | B0S3M8 | | 153 | 316 |
| 5 | A0A5R8Q9T8 | | 68 | 240 |
| 6 | V2XMF4 | N-term | 80 | 239 |
| | | C-term | 240 | 366 |
| 7 | R6V4J4 | | 37 | 420 |
| 8 | R3TX93 | | 40 | 276 |
| 9 | C6CUY3 | | 40 | 363 |
| 10 | A0A0F7RLJ7 | | 46 | 326 |
| 11 | A0A2V5K856 | Nterm | 38 | 201 |
| | | Cterm | 201 | 347 |
| 12 | A0A099WCN8 | | 48 | 417 |
| 13 | A0A069CUH0 | | 64 | 357 |
| 14 | A8MK03 | Nterm | 24 | 225 |
| | | Cterm | 225 | 337 |
| 15 | A0A4U7JL97 | | 37 | 420 |
| 16 | A0A494X6E7 | | 38 | 377 |
| 17 | A0A2Z4U801 | | 240 | 508 |
| 18 | R5FG57 | | 42 | 394 |
| 19 | S0JHJ4 | | 32 | 199 |
| 20 | A0A1I6IKX0 | Nterm | 39 | 142 |
| | | Middle | 143 | 244 |
| | | Cterm | 244 | 366 |
| 21 | A0A373LEL7 | Nterm | 30 | 196 |
| | | Cterm | 197 | 420 |
| 22 | R7HBU9 | Nterm | 38 | 158 |
| | | Middle | 158 | 251 |
| | | Cterm | 252 | 352 |
| 23 | Q81AL6 | | 21 | 339 |
| 24 | B1BZ86 | | 36 | 324 |

**Table S3: Structure model DALI results info:** The table lists the top DALI hits of the AlphaFold models for the N-terminal sequence clusters with the regarding Z-scores and similarity information when the structures are superposed using pymol. The Notes column gives information about the similarities to structures alternative to the top DALI hit, which were described in the main text.

| Nr. | Domain | Top DALI hit | Z-score | Superposition (RMSD) | Notes |
|---|---|---|---|---|---|
| 1 | | 5ny0:A | 27.3 | 5.07 Å over 272 res. | |
| 2 | N-term | 4b60:B | 11.5 | 3.42 Å over 128 res. | |
| | C-term | 4igb:B | 13.7 | 3.27 Å over 128 res. | 4jdz:A -> 3.74 Å over 136 res. |
| 3 | | 6poo:A | 10.5 | 3.24 Å over 64 res. | 4jdz:A -> 3.89 Å over 104 res. |
| 4 | | 2x9x:A | 7.5 | 4.43 Å over 112 res. | |
| 5 | | 5xcb:A | 10.8 | 4.52 Å over 128 res. | |
| 6 | N-term | 5cf3:A | 11.1 | 4.57 Å over 128 res. | |
| | C-term | 2z1p:A | 11.7 | 3.50 Å over 120 res. | |
| 7 | | 6kqs:A | 52.8 | 1.42 Å over 352 res. | |
| 8 | | 6fx6:A | 20.8 | 3.78 Å over 224 res. | |
| 9 | | 2z2o:B | 31.1 | 3.04 Å over 272 res. | |
| 10 | | 4nuh:A | 15.2 | 4.30 Å over 176 res. | |
| 11 | Nterm | 4nen:A | 11.5 | 4.06 Å over 120 res. | |
| | Cterm | 3rkp:A | 12.4 | 3.35 Å over 128 res. | |
| 12 | | 5ny0:A | 31.1 | 3.39 Å over 280 res. | |
| 13 | | 7nxd:A | 10.1 | 6.96 Å over 72 res. | 4igb:B -> 4.88 Å over 72 res. |
| 14 | Nterm | 6vxk:D | 8.1 | 7.37 Å over 120 res. | |
| | Cterm | 5l5g:D | 9.7 | 3.99 Å over 88 res. | |
| 15 | | 3bz5:A | 37.7 | 3.00 Å over 352 res. | |
| 16 | | 6v4p:A | 28.9 | 4.64 Å over 272 res. | |
| 17 | | 6fwv:B | 13.6 | 7.61 Å over 224 res. | |
| 18 | | 3n6z:A | 32.6 | 4.68 Å over 288 res. | |
| 19 | | 6fx6:A | 14.6 | 4.19 Å over 128 res. | 5A0L:B -> 4.26 Å over 144 res. |
| 20 | Nterm | 4uid:B | 9.1 | 3.94 Å over 72 res. | 2l04:A -> 2.92 Å over 64 res. |

| | | | | | |
|---|---|---|---|---|---|
| | Middle | 1f00:I | 8.7 | 4.19 Å over 80 res. | |
| | Cterm | 6k73:B | 6.8 | 5.48 Å over 80 res. | |
| 21 | Nterm | 6fx6:A | 11.7 | 4.22 Å over 120 res. | |
| | Cterm | 3kpt:A | 9.2 | 6.11 Å over 80 res. | |
| 22 | Nterm | 4u7k:G | 8.9 | 3.46 Å over 80 res. | |
| | Middle | 1yew:E | 7.6 | 4.24 Å over 80 res. | 5aea:A -> 3.69 Å over 88 res. |
| | Cterm | 4hu8:C | 8.8 | 3.99 Å over 72 res. | |
| 23 | | 3hjr:A | 8.7 | 4.46 Å over 128 res. | |
| 24 | | 6fwy:B | 13.4 | 5.35 Å over 232 res. | |

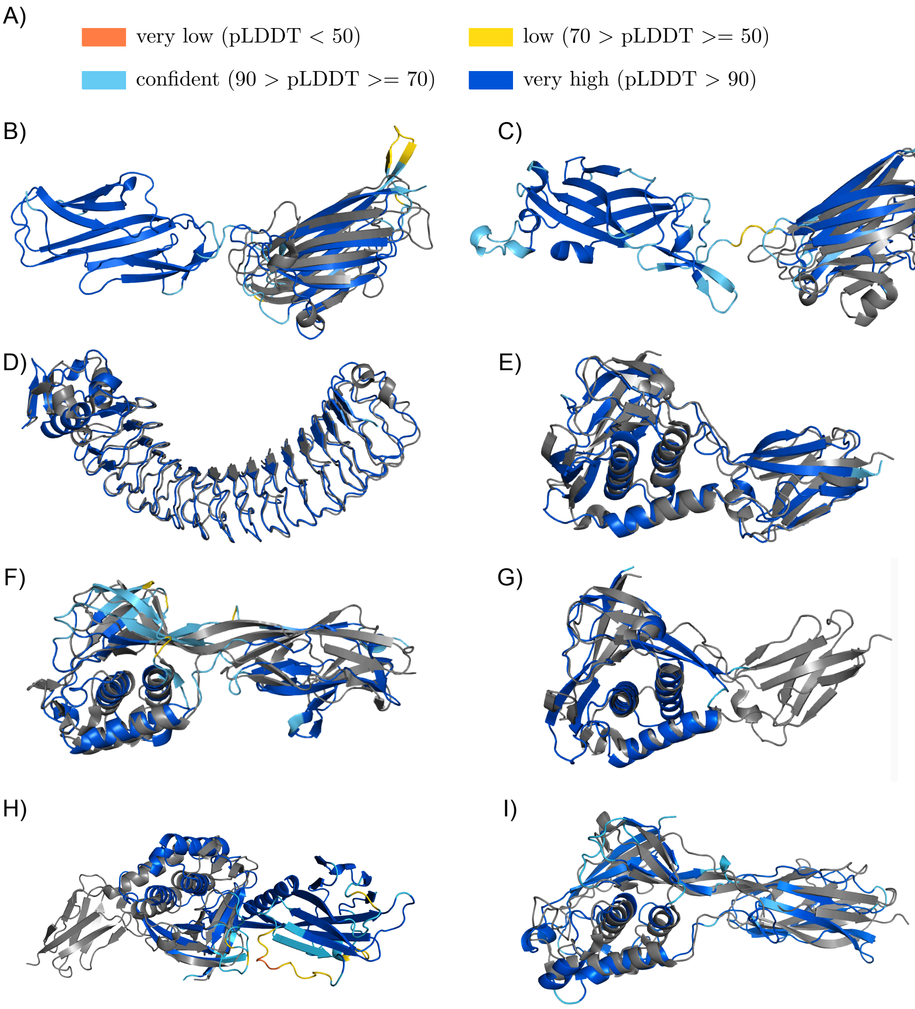


**Figure S2: Superposition of structure models shown in figure 5 with sequence similar adhesive domains found by Jackhmmer:** a) Colour legend representing the quality of the AlphaFold models. b) Superposition of the structure model of cluster group 2 with the SdrG_C_C binding domain (PDB:4jdz:A). c) Superposition of Collagen_bind domain (PDB:2z1p:A) to the C-terminal domain of the cluster group 6 structure model. d) Predicted structure model for cluster group 15 superposed with the functional region of Internalin J (PDB:3bz5:A). e) Predicted structure model for cluster group 8 superposed with S. aureus TED adhesive domain (PDB:6fx6:A). f) Predicted structure model of group 17 superposed with the Bacillus anthracis TED adhesive domain (PDB:6fwv:B). Superposition of S. aureus TED domain (PDB: 6fx6:A) and the structure model for g) cluster 19 and h) cluster 21. i) Cluster 24 structure model superposed to thioester domain of E. faecium (PDB: 6fwy-B).

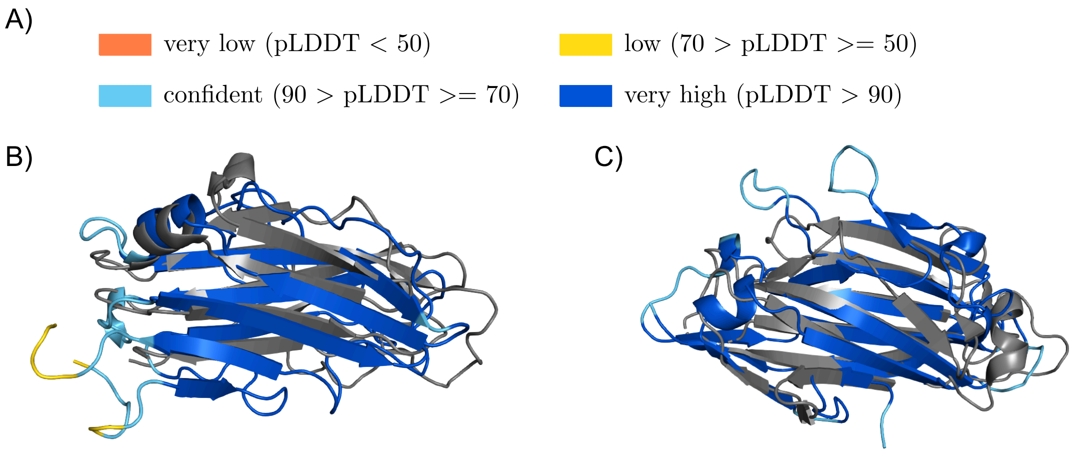


**Figure S3: Superposition with the DALI detected PDB structures similar to the clusters showing a jelly-roll like structure.** a) AlphaFold colour legend for the structure prediction confidence. b) Superposition of the structure model for cluster group 4 with the GramPos_pilinBB domain (PDB:2x9x:A). c) Predicted structure model for cluster group 5 superposed with the GramPos_pilinBB domain (PDB:5xcb:A).

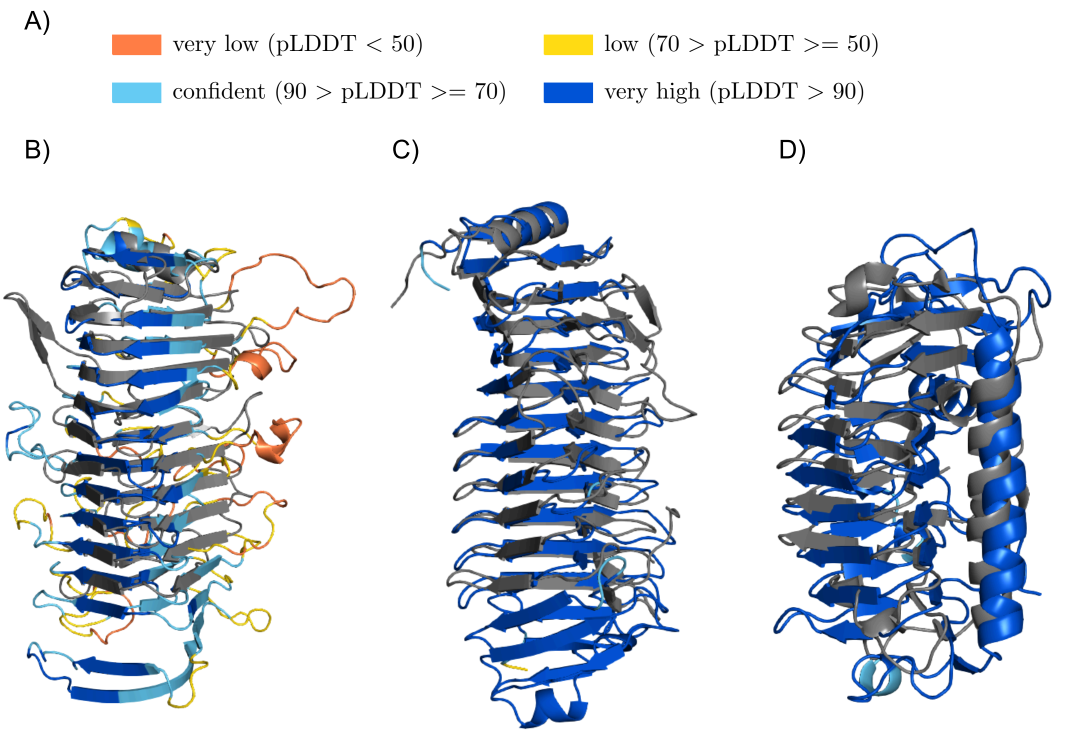


**Figure S4: Beta-solenoid predicted structure superposed with the top DALI matches in the PDB database.** a) AlphaFold prediction confidence colour legend. Superposition with the adhesive region of the serine-rich Lactobacillus reuteri adhesin (PDB:5ny0:A) with the b) predicted structure model for cluster group 1 and c) predicted structure for cluster group 12. d) Superposition of the structure model for cluster group 10 with the Ice_binding domain (PDB:4nuh:A).

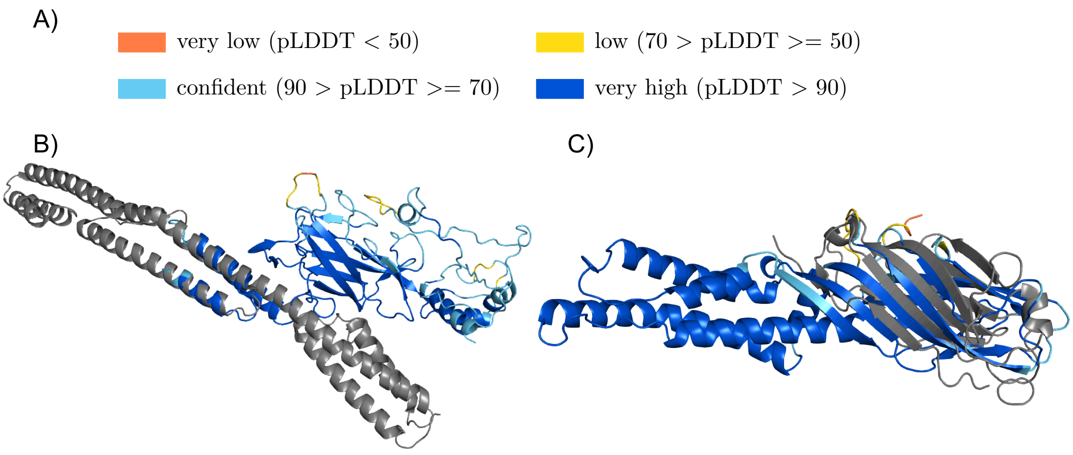


**Figure S5: Superposition of clusters with ambiguous adhesion function with DALI hits:** a) Confidence colour legend for AlphaFold predicted structures b) Cluster 3 structure model superposed with the immunogenic adhesin BibA (PDB:6poo:A). c) Cluster 13 structure model superposed with the S. gordonii Sgo0707_N2 domain (PDB:4igb:B).

**Table S4: Pfam families created based on this study**

| Cluster | Pfam accession | Pfam identifier |
|---|---|---|
| 1+12 | PF20585 | Pectate_lyase_5 |
| 4 | PF20592 | pAdhesive_10 |
| 5 | PF20595 | pAdhesive_6 |
| 10 | PF20597 | pAdhesive_15 |
| 13 | PF20609 | pAdhesive_17 |
| 24 | PF20610 | TED_2 |

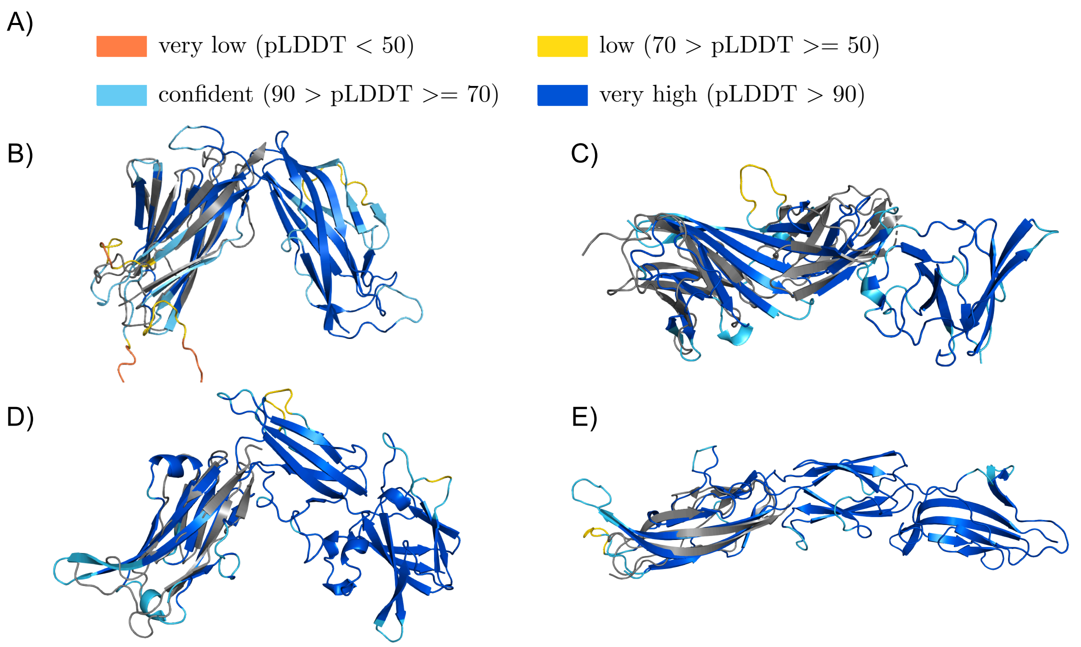


**Figure S6: Superposition of clusters with potential stalk characteristics with DALI hits:** a) Colour legend of AlphaFold prediction confidence b) Cluster 11 structure model superposed to stalk of an Integrin alpha-X protein (PDB: 4nen:A). c) Superposition of prediction structure for cluster 14 with the TIG domains of the plexin-C1 protein (PDB: 6vxk:D). d) Superposition of the N-terminal domain of the prediction structure for cluster 20 with Big_2 stalk domain of the major tail protein V (PDB: 2l04-A). e) Cluster 22 structure model superposed to the PKD_4 stalk domain of a collagenase ColH (PDB: 4u7k:G).

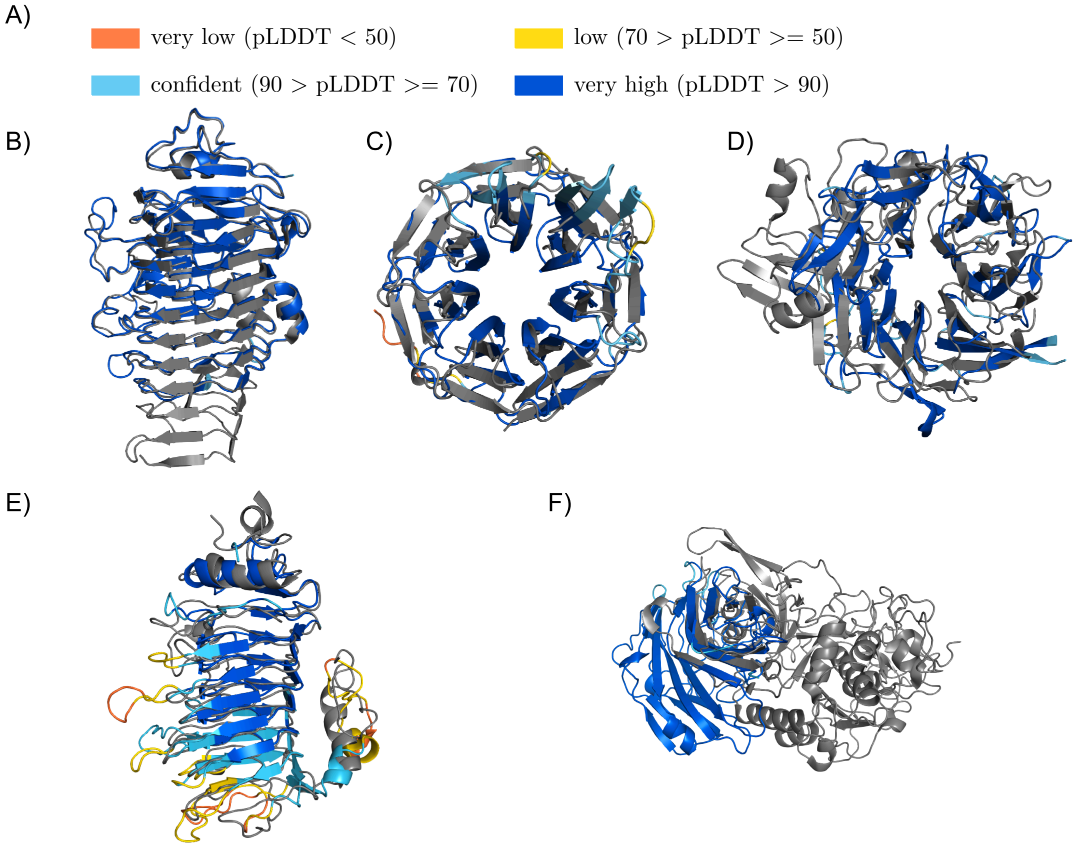


**Figure S7: Predicted structures for clusters with unlikely adhesion function:** (a) Colour legend for AlphaFold models quality. (b) Cluster 7 structure model superposed with an Eubacterium lacto-N-biosidase (PDB:6kqs:A). (c) Superposition of cluster 9 structure model with a virginiamycin B lyase (PDB:2z2o:B). (d) Cluster 16 structure model aligned with an human integrin alpha-IIb/beta-3, which binds among others to fibrinogen (PDB: 6v4p:A). (e) Superposition of cluster 18 structure model with a putative immunoglobulin protease (PDB: 3n6z:A), whereby the structure strongly resembles a pectate lyase. (f) Structure model of cluster 23 superposed to the P_proprotein domain of a protease (PDB: 3hjr:A).
